# Cross-individual translation of spontaneous zebrafish brain activity through a shared latent representation

**DOI:** 10.64898/2026.01.09.698719

**Authors:** Mattéo Dommanget-Kott, Jorge Fernandez-de-Cossio-Diaz, Guillaume Faye-Bédrin, Georges Debrégeas, Volker Bormuth

## Abstract

Spontaneous activity is a hallmark of brain function, reflecting the underlying circuit organization. Identifying conserved structure across individuals in this self-sustained activity has remained a longstanding challenge, especially in vertebrates where one-to-one neuron correspondence is inaccessible. Here, we introduce latent-aligned Restricted Boltzmann Machines (LaRBMs), an unsupervised generative approach that uncovers a common representational space from cell-resolved whole-brain recordings in larval zebrafish. This latent space consists of spatially localized co-activation motifs, or cell assemblies, that generalize across animals and form interpretable building blocks of population-wide activity. LaRBMs enable bidirectional mapping of instantaneous whole-brain activity patterns between individuals: activity patterns from one fish can be encoded into the latent space and decoded into another. The translated patterns are assigned high probability by the recipient model and retain the original spatial organization. These results show that spontaneous activity in the vertebrate brain is highly stereotyped at the level of functional cell assemblies and can be reliably captured through a common latent code. Because it provides an interpretable and quantitative framework for functional cross-individual alignment, LaRBM paves the way for comparative phenotyping of brain activity across developmental, genetic, and pathological variation.

**Significance Statement:** Spontaneous brain activity, without external stimuli, shapes development, constrains coding, and reflects neural organization. Whether this activity reveals similar organization across individuals has been unclear. Using single-cell, whole-brain recordings in larval zebrafish and statistical learning, we find that spontaneous dynamics share a common structure across animals. Spatially organized co-activated neuron assemblies recur across individuals and act as building blocks of population activity. This shared representation enables fictive translation of activity from one fish into another’s neural space while preserving spatial and statistical plausibility. These results suggest that brains organize population activity by similar principles to represent internal states. Our findings reveal conserved organization in the vertebrate brain and establish a quantitative framework for comparing brain dynamics across individuals and conditions.

## Introduction

Spontaneous brain activity is a fundamental feature of neural systems, shaping sensory processing, behavior, and internal state dynamics even in the absence of external stimuli [10, 23, 4]. In humans, population-level analyses of resting-state fMRI have revealed large-scale networks that are spatially and functionally conserved across individuals [9]. These so-called resting-state networks (RSNs) have been interpreted as a manifestation of intrinsic brain organization and are increasingly used to study brain development [40], aging [14], and psychiatric disorders [5]. However, their biological interpretability is limited by the low spatial and temporal resolution of fMRI, which makes it difficult to relate them to specific cell assemblies or circuit motifs.

At the other end of the spectrum, exhaustive recordings in small organisms, such as *C. elegans*, have enabled dynamic models that directly link activity to connectome structure. These models can predict transitions between behavioral states and reveal low-dimensional manifolds underlying brain-wide activity [3, 13, 2]. However, they are largely restricted to small nervous systems with stereotyped anatomy and limited neuron count.

In between these two extremes lies a major challenge: how to model spontaneous activity at single-cell resolution in vertebrates with conserved regional/mesoscale organization but without neuron-to-neuron correspondence, while retaining interpretability and enabling cross-individual comparisons. Zebrafish larvae offer a unique opportunity to fill this gap. Their optical transparency and compact brains enable functional imaging of nearly all neurons at single-cell resolution over extended periods [31, 1]. Recent studies have leveraged this animal model to study sensory-motor transformations [43, 26, 30], behavioral state dynamics [24, 29], spontaneous brain activity [41, 35, 33], or internal models [32].

Approaches to inter-individual comparison in neuroscience have mainly relied on anatomical registration combined with stimulus- or behavior-guided voxel-wise averaging or clustering. Although useful, these methods assume voxel-level correspondence and overlook the rich variabilities at neuronal scales. More recent frameworks such as Shared Response Models (SRMs) [7], hyperalignment [16, 15, 20, 21], spatial autoencoders [6], and contrastive frameworks for cross-subject alignment [36] aim to learn shared embeddings across subjects. However, most analyses remain limited to individual subjects or rely on anatomical alignment combined with stimulus- or behavior-guided averaging, making them unsuitable for extracting shared structure from spontaneous activity at the neuronal level.

Probabilistic generative energy-based models offer a powerful alternative [17]. By modeling the statistics of instantaneous neuronal configurations (a.k.a. *states*) **v**_*t*_ through a Boltzmann distribution *P*(**v**) ∝ *e*^−*E*(**v**)^, they have been used to model retinal activity, hippocampal replay [8, 34], and patterns of spontaneous activity [41, 27]. In particular, Restricted Boltzmann Machines (RBM) model the joint distribution of neural activity *P*(**v, h**) via latent variables (hidden units, *h*_*µ*_) that can be interpreted as co-activating neuronal assemblies. However, these studies have generally focused on single datasets, and it remains unclear whether RBMs can be used to identify conserved structures across individuals, especially when applied to full-brain recordings at cellular resolution.

Here, we introduce a framework for discovering a shared, interpretable latent structure in spontaneous brain-wide activity across individuals. Using whole-brain calcium imaging in larval zebrafish, we train RBMs with a shared latent space, a strategy we term Latent-aligned RBMs (LaRBMs), in which the hidden units are constrained to maintain their prior functional organization and activation distribution across individuals. This enables us to extract a common set of latent motifs, corresponding to spatially organized assemblies, that are reused across different brains. Crucially, we go beyond statistical comparison by encoding activity from one fish into the shared latent space through *P*(**h** | **v**) and decoding the resulting latent representation through *P*(**v** | **h**) into the neural space of another fish. We show that spontaneous patterns at single-cell resolution can be mapped across individuals, while preserving both spatial organization and statistical structure.

Our approach fills a key gap between connectome-informed modeling in C. elegans and population-level RSN analysis in humans. It enables comparative, interpretable, and probabilistically grounded modeling of spontaneous brain-wide activity across individuals. This provides a concrete route for identifying shared circuit motifs that shape spontaneous dynamics at the mesoscale, bringing us closer to a common representational framework for a vertebrate brain.

## Results

### Latent representations of spontaneous brain-wide activity are variable and non-alignable across individual RBMs

We showed previously in van der Plas et al. [41] that spontaneous whole-brain neuronal activity of zebrafish larvae could be modeled by Restricted Boltzmann Machines (RBMs). In this framework, the binarized spiking activity of individual neurons is mapped onto a latent space comprised of hidden units that correspond to cell assemblies (*i.e*. functional ensembles of co-activating neurons; Fig. 1 A). Each hidden unit *µ* is characterized by the spatial distribution of weights *w*_*iµ*_, where *i* is the neuronal index. These weights are found to be sparse, spatially compact, and define functional circuits and anatomical structures. This biological interpretability raises the appealing possibility that RBMs trained on different fish might yield comparable latent assemblies that could support cross-individual comparisons. Whether such convergence occurs in practice, however, is not obvious given the stochastic nature of RBM training and potential degeneracy of the learned representations. We therefore start by testing the stability and alignment of RBM latent features across independent trainings.

**Fig 1.**
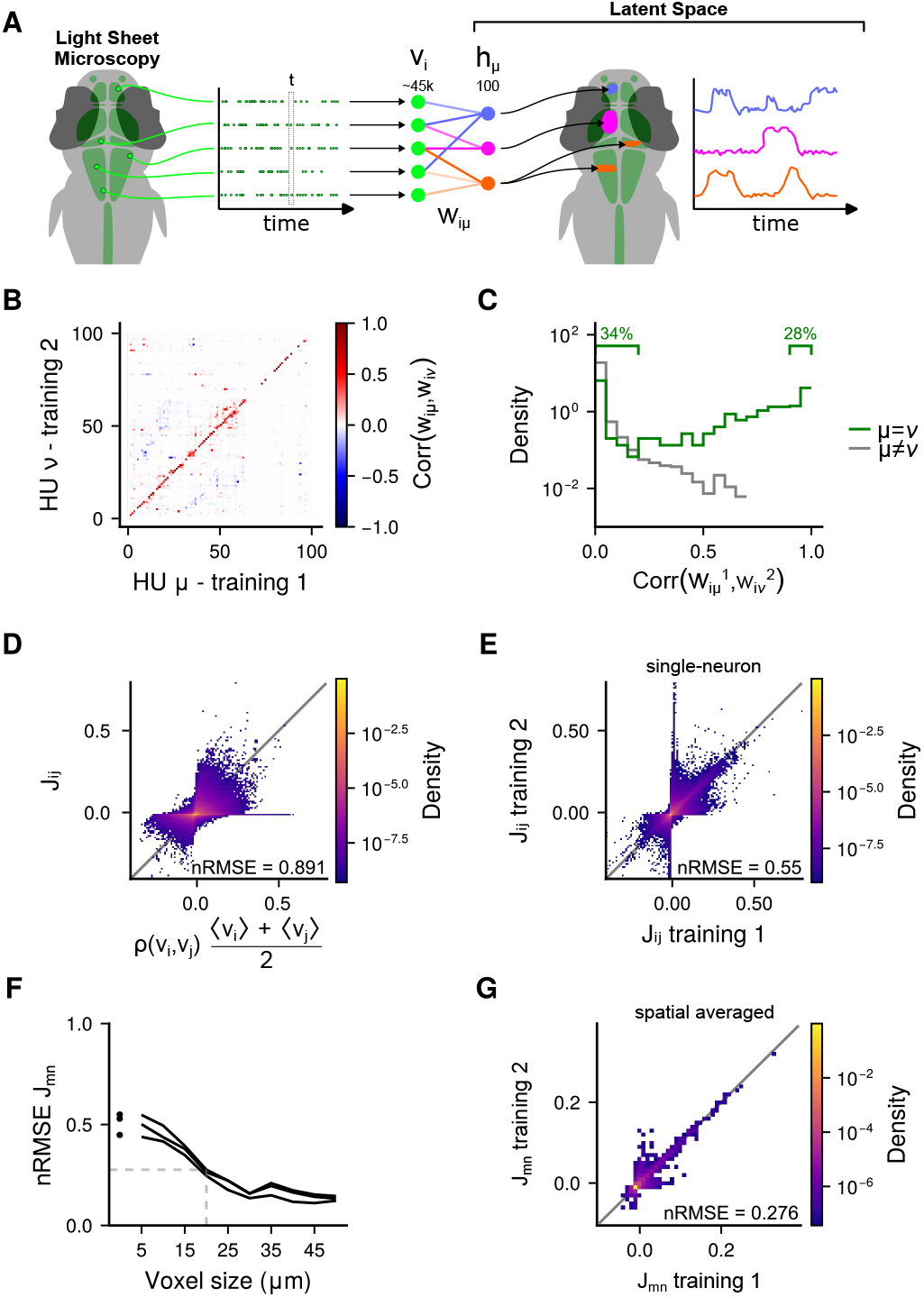
Restricted Boltzmann Machines (RBMs) produce degenerate representations of neuronal functional organization. **A**: Zebrafish larvae neuronal activity is recorded using Light Sheet Fluorescence Microscopy and deconvolved into binarized spike trains. RBMs are trained with *N* ≈ 45000 binary visible units *v*_*i*_ corresponding to neurons and connected to *M* = 100 dReLU hidden units *h*_*µ*_ via a weight matrix *w*_*iµ*_ (see Methods Restricted Boltzmann Machines). The hidden layer describes a latent space capturing the activation modes of the whole neuronal population as groups of co-activating neurons [41]. **B**: Pairwise Pearson correlation between the weight matrices *w*_*iµ*_ and 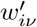 of two example RBMs trained on the same neuronal recording. The matrix rows have been reordered to obtain the best alignment between the two trainings. **C**: Distribution of correlations from panel B, for best alignment pairs (*ν* = *µ*, green) and other pairs (*ν*≠ *µ*, grey). **D**: Rescaled pairwise Pearson correlation 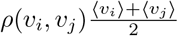 between neurons *i* and *j* vs. coupling matrix *J*_*ij*_ (see Eq.14) inferred from an example RBM. **E**: Comparison of two example coupling matrices inferred from two RBMs trained on the same neuronal recording. **F**: nRMSE (see Methods Measuring identity) between coupling matrices inferred from three different training on the same neuronal recording. Dots correspond to *J*_*ij*_ matrices as presented in panel C. Lines correspond to coupling matrices spatially averaged on a cubic-voxel grid : 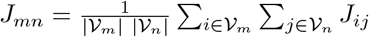 with *V*_*m*_ and *V*_*n*_ the sets of neurons in voxels *m* and *n* respectively. **G**: Comparison of two example coupling matrices *J*_*mn*_ spatially-averaged at 20*µm* voxel size.

### RBMs represent neuronal statistics from a degenerate set of latent features

To assess the variability in representation we investigated the robustness of the latent features by training multiple RBMs on the same neuronal dataset (fish 1 in Tab. 1). We used the same training parameters and selected the models that had reached a high level of statistical performance (see Methods Teacher RBMs and Evaluation).

**Table 1.**
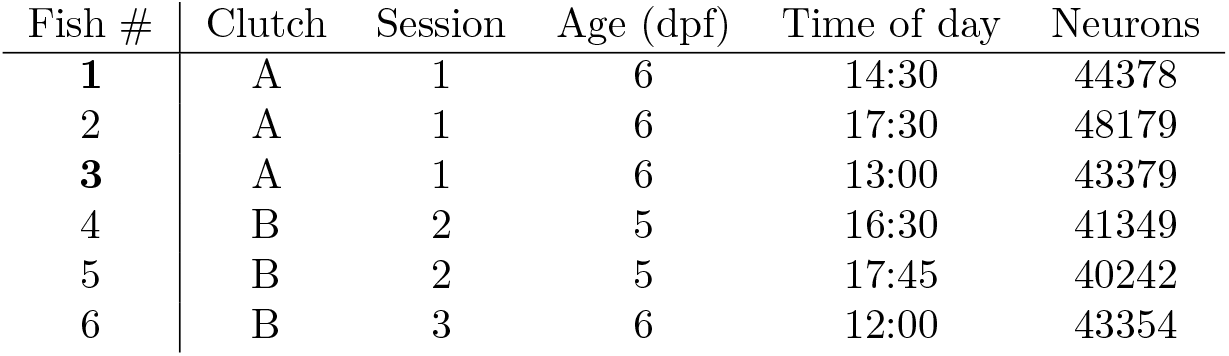
List of fish.

| Fish # | Clutch | Session | Age (dpf) | Time of day | Neurons |
| --- | --- | --- | --- | --- | --- |
| 1 | A | 1 | 6 | 14:30 | 44378 |
| 2 | A | 1 | 6 | 17:30 | 48179 |
| 3 | A | 1 | 6 | 13:00 | 43379 |
| 4 | B | 2 | 5 | 16:30 | 41349 |
| 5 | B | 2 | 5 | 17:45 | 40242 |
| 6 | B | 3 | 6 | 12:00 | 43354 |

We then computed the Pearson correlation 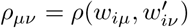 between the weight matrices *w*_*iµ*_ and 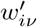 of two trainings. For the best alignment, we found that ≈ 28% of hidden units (out of 100) could be unequivocally paired between trainings (*ρ*_*µν*_ *>* 0.9), while ≈ 34% had no matching pair (*ρ*_*µν*_ *<* 0.2) (see Fig. 1 B-C). This indicates that only a core set of hidden units are stable across trainings.

We performed the same analysis on the expected hidden activity *h*_*µ*_(*t*) = 𝔼[*h*_*µ*_ | **v**(*t*)] by computing the correlation *ρ*_*µν*_ = *ρ*(*h*_*µ*_(*t*), *h*_*ν*_ (*t*)), and found that ≈ 54% of hidden units had correlations *ρ*_*µν*_ *>* 0.9 and ≈ 18% *ρ*_*µν*_ *<* 0.2 (see Fig. S1). This result reflects the fact that, despite being connected to different neuronal populations, many hidden units share similar temporal signals, capturing dominant dynamics of brain activity spanning large brain regions.

### RBMs produce stable functional connectivity model only at coarse-grained scales

We showed previously [41] that effective couplings *J*_*ij*_ between neurons *i* and *j* could be inferred from the model (see Sec. Inferred neuronal couplings), and that this measure of functional connectivity reflects, at least in part, the structural connectivity measured between brain regions. Importantly, *J*_*ij*_ is not a measurement of the correlation between neurons, but reflects a model of connectivity which takes into account indirect network interactions (see Fig. 1 D).

We compared the effective couplings *J*_*ij*_ inferred from multiple trainings on the same dataset as described above. We found that, while they were strongly correlated, couplings were also highly variable, with a significant fraction of neuron pairs being strongly coupled in one model and completely decoupled in another (Fig. 1 E). However, when couplings were spatially averaged over cubic-voxels of side ⪆ 20*µm*, the resulting *J*_*ij*_ was much more stereotypical between trainings (Fig. 1 F-G). This suggests that, while the inferred couplings between neurons are degenerate between RBM trainings, they are well defined at a coarse-grained scale.

In summary, we found that RBMs provide partially degenerate representations of neuronal activity, reflecting the stochastic nature of the model and its training algorithm, as well as insufficient constraints from the data. These findings caution against naively comparing RBMs trained on different individuals, particularly at neuronal scales. However, the stability of the model at mesoscopic scales suggests that RBMs could capture a stereotypical organization of spontaneous brain activity which might be comparable across individuals, motivating multi-individual training paradigms that explicitly align latent spaces.

### A global voxel-level RBM uncovers shared structure in spontaneous brain activity across individuals

We first asked whether spontaneous whole-brain activity across individual zebrafish larvae could be described by a common latent representation, at a coarse-grain level, using anatomical cross-individual registration.

To enable cross-individual comparisons, we spatially registered six fish to a common brain coordinate system using affine and non-linear transformations (see Methods Morphological Registration). The resulting aligned spontaneous brain activity from all six fish is visualized side by side in Movie S1, with each fish shown separately to highlight individual dynamics within the shared coordinate space. Each neuron was then assigned to a voxel in a 3D grid (20 *µm* cube size), and voxel activity was computed as the mean of the normalized fluorescence (Δ*F/F* ) of all neurons within it (Fig. 2 A and Methods Voxelisation). Only voxels that contained at least two neurons in each fish were retained, ensuring that anatomical coverage was consistent across individuals. This filtering step yielded a total of 2,995 shared voxels, with an average of 11 *±* 5 neurons per voxel (Fig. S2 D mean and standard deviation). Importantly, this voxel size was the smallest possible to retain *>* 75% of all neurons in all fish, thus balancing spatial resolution with inter-individual consistency (Fig. S2 B).

**Fig 2.**
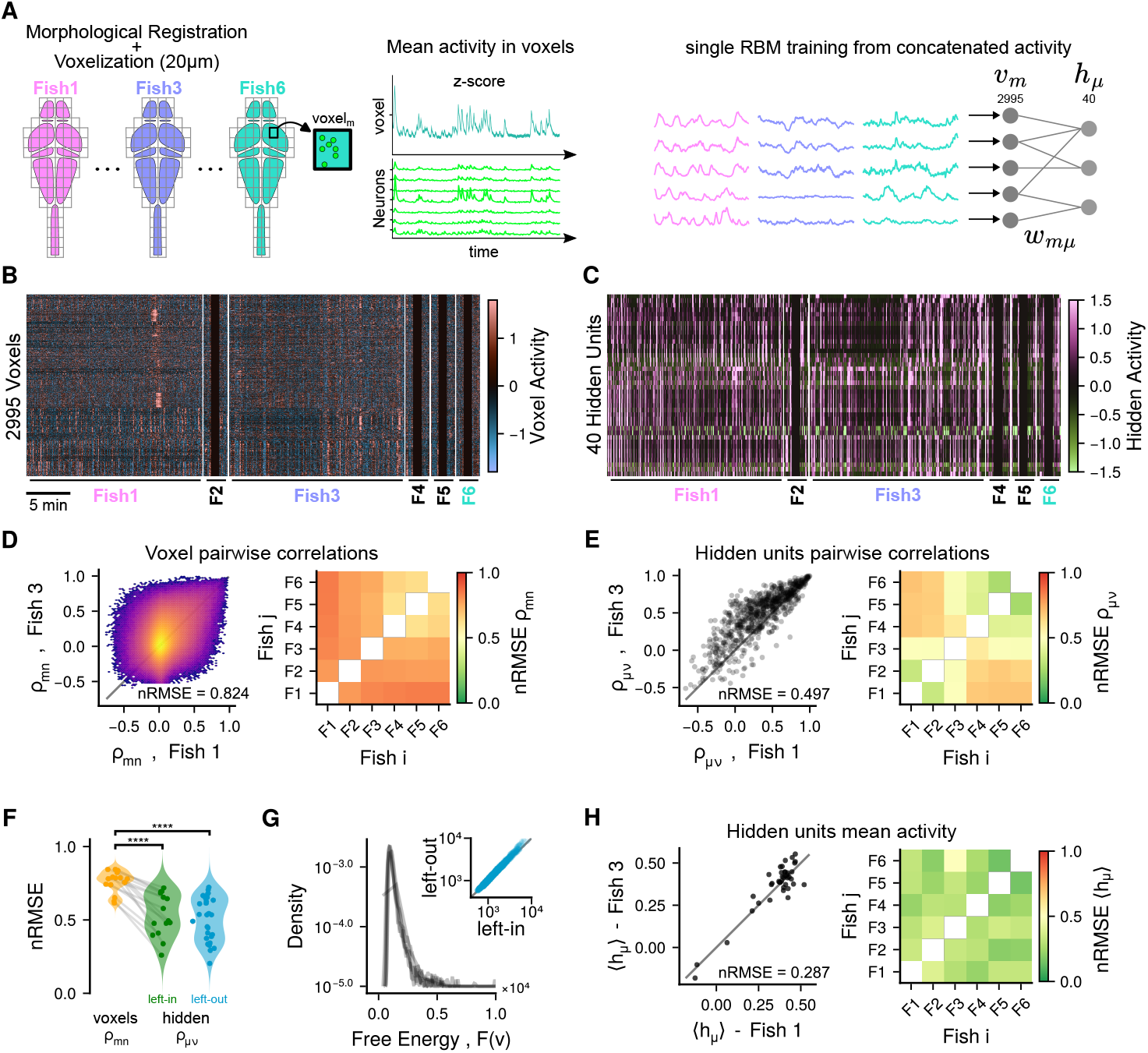
Concatenation and voxelization allow for a common RBM representation. **A**: Pipeline of voxelized RBM training. Averaged brain scans are first morphologically registered to a common reference brain. A grid of cubic voxels with side 20*µm* is then used to group neurons spatially into a total of *N* = 2995 voxels. The neuronal activity within each voxel is averaged and z-scored. The voxel activity for all 6 fish is concatenated to construct a single dataset, which is then used to train an RBM (see Methods Voxelised RBMs). **B**: Concatenated voxelized activity of all 6 fish. Fish 1 and 3 are shown in full, while the activity of fish 2, 4, 5 and 6 are truncated for visualization purposes. **C**: The activity of panel B *translated* into the hidden space of the RBM (*M* = 40 hidden units, **h**(*t*) = 𝔼_**h**|**v**_[**v**(*t*)], see Methods Translating neuronal activity patterns across individuals). **D**: Stereotypy of the pairwise Person correlation coefficient *ρ*_*mn*_ between voxels *m* and *n*. Left: example fish pair. Right: nRMSE (see Methods Measuring identity) of *ρ*_*mn*_ between all pairs of fish. **E**: Stereotypy of the pairwise Person correlation coefficient *ρ*_*µν*_ between hidden units *µ* and *ν*. Left: example fish pair. Right: nRMSE of *ρ*_*µν*_ between all pairs of fish. **F**: Distributions of fish-to-fish nRMSE for pairwise Pearson correlations between voxels (left orange, same as panel D right) and between hidden units (middle and right, respectively green and blue). The middle distribution (left-in, green) is obtained from a single RBM trained on all six fish combined (same as panel E right). The right distribution (left-out, blue) is derived from comparisons between a fish excluded from training and each of the five fish that were included in training, with all possible pairs of trained fish and left-out fish pooled into this distribution. Each dot represents a pair of fish. The p-value was computed with a one-tailed Mann-Whitney U test (∗∗∗∗ means *p <* 0.0001). **G**: Distribution of free energy *F*(**v**) (see Methods Restricted Boltzmann Machines) of voxel activity configurations **v**. One line per fish. Inset: Comparison of free energy distributions of an example fish when included (left-in) or excluded (left-out) from the training. **H**: Stereotypy of the mean activity 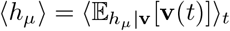 of hidden units *µ*. Left: example fish pair, one point per hidden unit. Right: nRMSE of ⟨*h*_*µ*_⟩ between all pairs of fish.

We then trained an RBM on the concatenated voxelized activity from all fish (Fig. 2 A-B). This global RBM consisted of *N* = 2995 Gaussian visible units (corresponding to the number of shared voxels) and *M* = 40 hidden units (see Methods Training). Before training, we *z*-scored each voxel’s activity within each fish to account for differences in mean and variance across individuals (see Methods Voxelisation) [19]. Without this normalization, voxelized training resulted in the RBM capturing fish identity rather than shared features.

### The global RBM is a good model for data from individual fish

Validation of this global RBM model on each fish separately demonstrated that the model accurately captured pairwise voxel correlations in 5 out of 6 fish, and more generally, provided a good statistical model of voxelized activity across individuals (see Fig. S4). This indicates that, despite voxel-level variability, the RBM can extract a common statistical structure. This is particularly notable because no fish-specific information was provided during training.

### Latent representations are more stereotyped than raw activity

Projecting each fish’s activity into the RBM’s hidden space (Fig. 2 B-C) revealed that correlations between hidden units were more conserved across individuals than correlations between voxels (Fig. 2 D-F). This was quantified using the normalized Root Mean Squared Error (nRMSE). Pairwise voxel correlations had high fish-to-fish variability (nRMSE up to ∼ 0.8) with weak voxel size dependence (Fig. S2 G). In contrast, hidden-unit correlations were significantly more stereotyped (nRMSE ∼ 0.3–0.7, Mann–Whitney U test *p<* 0.01). This remains true even when a single fish is excluded from the training (Fig. 2 F), confirming that the global RBM captures the inter-individual regularities present in the data and does not merely learn an *ad hoc* and over-fitted representation space which matches all training individuals. Interestingly, we observed that fish from the same clutch (F1–F3 vs. F4–F6) clustered together in the latent representations (see Fig. 2 E), although no explicit labeling was provided. This suggests that the RBM captures subtle commonalities in spontaneous activity linked to biological or experimental context. However, it is difficult to pinpoint the potential causes of these observed differences between clutches. The preparation and recording of the fish followed the same protocol, the imaging and registration quality were comparable across all fish, and the two clutches were recorded two weeks apart on the same microscope. We hypothesize that the potential differences between clutches could be due to a combination of the following factors: genetic differences (different sets of parents), temperature (which was not controlled during the experiment), and experimenter-related differences during fish handling.

### RBM captures shared structure without encoding individual idiosyncrasies

To verify that the model did not simply encode for fish-specific features, we considered evaluating the probability *P*(**v**) of neuronal configurations. Evaluating *P*(**v**) is computationally impractical, and we used the free energy as an alternative:

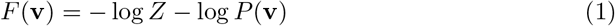

where *Z* is the partition function (see Methods Definition). We found that free energy distributions for all fish overlapped closely (Fig. 2 G), indicating that each individual’s activity was equally well-explained by the model. Once again, this remains true for fish excluded from training, showing that the model does not learn an over-fitted representation.

We further compared the mean hidden activity ⟨*h*_*µ*_⟩ across fish, and found it to be comparable between individuals (Fig. 2 H), suggesting that the RBM did not merely classify individuals.

Overall, these findings confirmed that the latent space was shared and not specialized to any specific fish. The dynamics of activity in this space might be different for different fish, but the overall distribution is similar. As such, it describes the common realm of possible hidden configurations **h**.

Despite anatomical variability, spontaneous brain activity across zebrafish larvae can be represented in a shared latent space. A single RBM trained on voxelized and concatenated data successfully captures second-order activity statistics across individuals. Importantly, latent representations show greater inter-fish stereotypy than voxelized activity, revealing higher-order regularities in collective dynamics. Crucially, the increased inter-fish stereotypy in hidden space cannot be attributed solely to compressing 2995 voxels into 40 latent variables. To test this, we compared fish-to-fish similarity of second-order covariance structure in voxel activity and in inferred hidden-unit activity (see Fig. S3 D). Hidden-space covariance structure is consistently more aligned across individuals than voxel-space covariance structure, indicating that the RBM performs a non-linear re-encoding into conserved co-activation motifs, rather than a generic low-dimensional projection. These findings establish the feasibility of constructing unified and interpretable representations of spontaneous neural activity across individuals.

### Single-cell resolved LaRBMs reveal conserved spatial cell assemblies via a shared latent space

While voxel-based registration enabled us to construct a common latent space at a coarse spatial resolution, this approach sacrifices single-neuron interpretability. Next, we asked whether we could preserve interpretability while enforcing a shared representation of spontaneous whole-brain activity across individual fish.

We developed a training framework in which a reference RBM, the *teacher*, defines the latent space, and subsequent *student* RBMs are trained on different fish under the constraints that they use the same latent space (Fig. 3 A). Each RBM models the binarized spiking activity of 43480 *±* 2755 neurons (mean and std, see Materials List of fish) recorded in individual fish at single-cell resolution.

**Fig 3.**
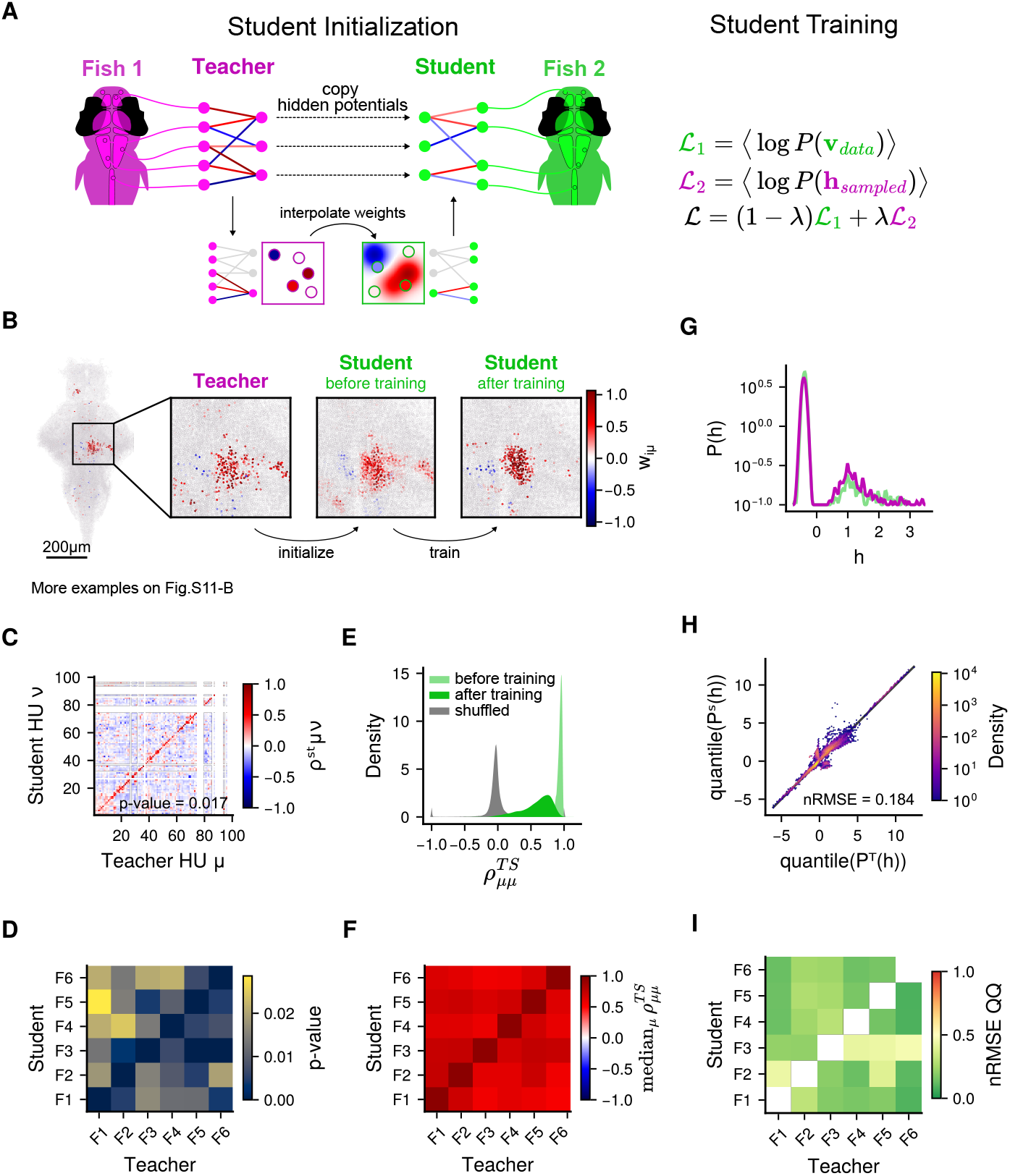
RBMs can be trained from priors and constrained to enforce a similar hidden space. **A**: Teacher/Student paradigm. A *student* LaRBM is initialized from a pretrained *teacher* RBM by copying its hidden layer and spatial interpolation of its weights. The student RBM is then trained on both student neuronal data and hidden configurations sampled from the teacher RBM. **B**: Example spatial interpolation of weights. Left: z-projection map and inset of teacher weights 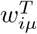 for each neuron *i* and a single hidden unit *µ* (out of *M* = 100). Middle: interpolated weights in student. Right: weights after student training. **C**: Example spatial correlation matrix 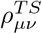 (see Methods Measuring Spatial Similarity) between weights maps of hidden unit *µ* of teacher RBM and hidden unit *ν* of student RBM. **D**: P-value of diagonal dominance of the 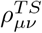 matrix (see Methods Measuring Diagonal Dominance) for every pair of teacher and student. **E**: Distribution of spatial correlations 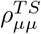 between corresponding hidden units in all teacher-student pairs for students before training (light green), students after training (green), and shuffled pairs of hidden units (grey). **F**: Median of spatial correlation median_*µ*_ 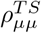 for every teacher-student pair. **G**: Prior distribution of hidden value *P*(*h*) for the example hidden unit presented in panel B, in both teacher (magenta) and trained student (green) RBMs. **H**: Quantile-quantile (Q-Q) plot of *P*(*h*) in the example teacher and student pair for all hidden units combined. **I**: nRMSE (see Methods Measuring identity) of Q-Q plot in panel H for every teacher-student pair.

In practice, we first trained the teacher RBM on one fish, with binary visible units and *M* = 100 hidden units to define the latent space (Methods Teacher RBMs and Supplementary Hyper-parameters selection for a more detailed discussion of the choice of hyperparameters). Each hidden unit *µ* in this trained teacher model represents a distributed functional assembly, a spatial pattern of coactive neurons, as defined by its weight vector *w*_*iµ*_ (see examples in Fig. 3 B and Fig. S7 B). Student RBMs have the same number of hidden units, but can have a different number of visible units to account for variability in neuron-count per fish. To initialize the student RBM parameters, we interpolated the spatial weight patterns from the trained teacher onto the neuron positions in the student brains, we copied the hidden unit potentials 𝒰_*µ*_(*h*) of the teacher, and we estimated the fields of the visible units from the corresponding neuron’s baseline activity in the student (see Methods Student LaRBM Initialization). These pre-initialized student RBMs do not capture the statistical structure of their respective fish, as they have not yet been trained on the new data. If trained without additional constraints, the student representations would adapt solely to their local data and gradually drift away from the teacher’s latent structure. We therefore introduced a new paradigm, which we call latent-aligned RBMs (LaRBMs), in which the student RBM is trained not only to maximize the likelihood of its own observed neuronal activity statistics, but also to capture the distribution of hidden-unit activations sampled from the teacher RBM:

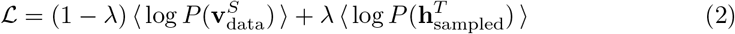

where 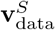 is the neuronal activity of the student, 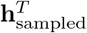 are hidden configurations sampled from the teacher RBM, and *λ* ∈ [0, 1] is an hyperparameter controlling the relative impact of each term on training (see Methods Latent-aligned Restricted Boltzmann machines (LaRBMs) for details and Fig. 3 A). This approach ensures that all student RBMs share a subspace of hidden configurations aligned with the teacher, while retaining sufficient flexibility to adapt to individual neuronal datasets.

### LaRBM students converge faster and more reliably

The LaRBM students were found to converge rapidly, requiring only one-tenth the number of training steps compared to unconstrained and randomly initialized models. This suggests that the initialization step acts as pretraining, while the actual training merely fine-tunes the model to match student statistics. Indeed, we found that, contrary to teachers where multiple RBMs needed to be trained to obtain a good model, student LaRBMs were consistently able to reproduce the first- and second-order statistics of neuronal activity, sometimes even exceeding that of the teachers (see Fig. S5 A-B and Fig. S6). Despite being trained with a latent-space constraint, student RBMs remain faithful to the distribution of neuron firing rates and pairwise covariances in their respective datasets. Notably, even though training does not explicitly account for correlations between hidden units, we found that their pairwise covariances were partially preserved between the teacher and student (see Fig. S5 C-D). This consistency across animals supports the hypothesis that the RBM’s hidden units reflect stereotyped functional connectivity motifs.

### Hidden units retain their spatially localized patterns

Each hidden unit in the teacher RBM represents a spatially localized pattern of co-activating neurons. After interpolating the teacher weights into a student RBM and training the model, we observed that these spatial patterns were largely retained (examples in Fig. 3 B and Fig. S7 B). The spatial correlation between corresponding hidden units in the teacher and student RBMs was on average above 0.6 and significantly higher than between non-matching pairs (see Fig. 3 C-E, bootstrap p-value ≈ 0.02, see Methods Measuring Spatial Similarity and Measuring Diagonal Dominance), indicating that most hidden units can be reliably matched between models based on their spatial weight patterns. We observed that student weight maps were generally of lower amplitude and more compact, that is, involving fewer strongly connected neurons 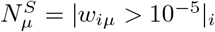 with 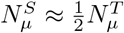 (see Fig. S7 A-B). This is consistent with the new likelihood at Eq. 2 and the weight regularization applied during training (see Supplementary Section Small weight expansion of the LaRBM). Crucially, the core spatial identity of the assemblies was maintained.

### Student RBMs preserve the teacher’s hidden-unit priors

Finally, we examined whether student RBMs accessed the same regions of the latent space as the teacher. We compared the prior distributions *P*(*h*_*µ*_) of each hidden unit and found close agreement between the teacher and student RBMs (Fig. 3 G-I). Quantile–quantile (Q–Q) plots showed strong linear correspondence (Fig. 3 H), and the identity error (nRMSE) across all teacher-student pairs was low (Fig. 3 I). This indicates that, despite anatomical and statistical variability, spontaneous activity across individuals can be mapped to the same latent sub-space.

Together, these results demonstrate that RBMs trained on different individuals can share a common latent space in which functional assemblies are reliably matched based on spatial structure and prior activation. However, this alone does not guarantee that individual brain states (*i.e*. single-time-point whole-brain spontaneous activity patterns) are encoded similarly across fish.

### Cross-individual translation confirms conserved encoding of spontaneous activity in the shared latent space

Restricted Boltzmann Machines enable encoding of neuronal configurations into the latent space, and decoding of latent configurations back into the neuronal space. This bidirectional generative capacity, combined with our shared latent representations, enables the translation of activity pattern from one individual to another through the latent space. Here, we evaluate whether such synthetic (translated) activity patterns share the same characteristics as the endogenous brain activity.

Specifically, starting from a neuronal configuration **v**^*T*^ from a teacher fish, we successively computed the expected hidden activity 𝔼[*h*^*T*^ | **v**^*T*^ ] and the decoded activity pattern 𝔼[**v**^*S*^ | *h*^*T*^ ] in the student brain (see Methods Translating neuronal activity patterns across individuals). We then assessed whether this translated activity is highly probable under the student model and retains the same spatial structure as the original teacher pattern. Because the mapping is bidirectional, it allows us to benchmark both teacher-to-student (*T* → *S*) and student-to-teacher (*S* → *T* ) translations against within-fish reconstructions (*T* → *T* or *S* → *S*) and randomized controls (Fig. 4 A). We show in Movie S3 and Movie S4 two example movies of **v**^*T* →*S*^ and **v**^*S*→*T*^, respectively, to illustrate this activity translation. Qualitatively, these movies show that this procedure is indeed capable of generating activity which resembles real brain activity, with translated patterns recapitulating salient features of spontaneous brain-wide activity, including spatially organized assemblies and bilateral coordination.

**Fig 4.**
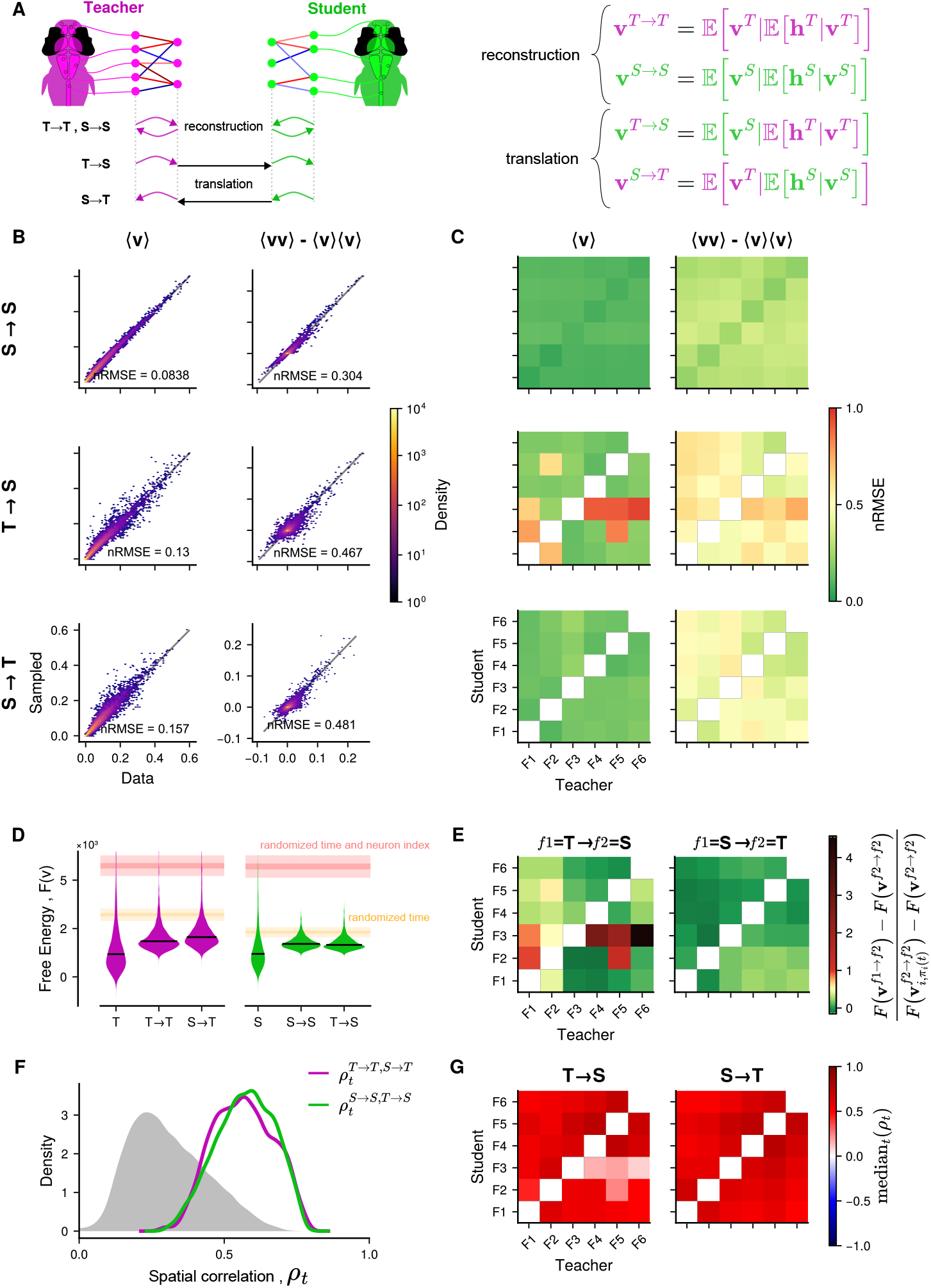
Spontaneous neuronal activity can be translated through the latent space to another fish. **A**: Neuronal configurations can be represented in hidden space (*M* = 100 hidden units) and then either reconstructed in the original neuronal space, or translated to another neuronal space. (see Methods Translating neuronal activity patterns across individuals) **B**: Neuronal statistics for the example teacher-student pair. Temporal average neuronal activity ⟨*v*_*i*_⟩_*t*_ (left) and pairwise covariance ⟨*v*_*i*_*v*_*j*_⟩ _*t*_ − ⟨*v*_*i*_⟩ _*t*_ ⟨*v*_*j*_⟩ _*t*_ (right) between empirical data and reconstructed student configurations (*S* → *S*, top) and translated configurations in both teacher-to-student (*T* → *S*, middle) and student-to-teacher (*S* → *T*, bottom). **C**: nRMSE (see Methods Measuring identity) for each panel of B and for every teacher-student pair. **D**: Distributions of free energy for empirical (*T* or *S*), reconstructed (*T* → *T* or *S* → *S*), and translated (*S* → *T* or *T* → *S*) configurations in an example teacher (left) and student (right) RBMs. Black lines indicate median. Horizontal bands indicate 50% and 99% of free energy distribution for randomized data with shuffled time frames (orange), and shuffled time frames and neurons (red). **E**: Median free energy contrast 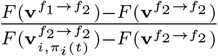 of translated configurations 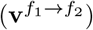 with respect to reconstructed configurations 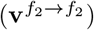 and time-randomized configurations 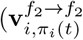, horizontal orange in panel D). Left panel: *f*_1_ =teacher and *f*_2_ =student. Right panel: *f*_1_ =student and *f*_2_ =teacher. Values around zero (green, 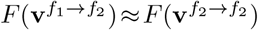) correspond to fish pairs where translated and reconstructed free energy have similar median values, while values *>* 1 (red, 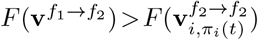) correspond to fish pairs where translated free energy is higher (*i.e*. lower probability) than a time-randomized bootstrap. **F**: Spatial correlation 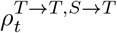 (magenta) and 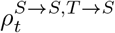 (green) between reconstructed and translated neuronal configuration for each time frame of an example teacher-student pair. In gray, the distribution of spatial correlations between shuffled pairs of time frames. **G**: Median of the distributions of spatial correlations from panel F for every teacher-student pair.

### Statistical features are preserved during mapping

We first compared the mean activity ⟨*v*⟩ _*t*_ and pairwise covariances ⟨*vv*⟩ of translated versus actual neuronal configurations. In both directions and for all pairs (*T* → *S* and *S* → *T* ), translated configurations preserve mean activity with high fidelity, and captured substantial pairwise covariances, except for a few outliers discussed below (see Fig. 4 B-C). These values were comparable to, and in some cases better than, within-animal reconstructions, indicating that translation via the latent space preserved meaningful structure. A detailed examination of the outliers reveals that the model fails to reproduce the mean activity ⟨*v*_*j*_⟩ of 2% to 18% of their neurons (see Fig. S9 A-B). These neurons tend to be spatially structured like functional ensembles (see Fig. S9 C), and we show in Fig. S9 D that this is consistently due to *<* 5 hidden units (out of 100), which apparently failed to maintain the teacher’s spatial distribution. To test whether the outliers arose from a convergence failure during RBM training, we retrained two of the outlier student RBMs. In both cases, it was possible to obtain models without failing hidden units, and the previously failing hidden units recovered the spatial organization of the corresponding teacher hidden units, removing this problem. However, these examples illustrate how a few hidden units can have a significant detrimental impact, confirming the robustness of the method the rest of the time.

### Translated patterns are probable under the receiving model

To test whether translated configurations were plausible under the receiving RBM, we evaluated their free energy *F*(**v**), which measures the log-probability of the configurations up to an additive constant. Translated patterns (*T* → *S* and *S* → *T* ) had free energies comparable to those of within-fish reconstructions (*T* → *T, S* → *S*), and well within the expected distribution for each fish (Fig. 4 D). In contrast, randomized activity patterns yield, under the same reconstruction procedure, significantly higher free energy and thus are unlikely under the receiving model. This latter test demonstrates that the translation accuracy is not a simple byproduct of the mapping procedure, but reflects the accuracy of the representation in hidden space. We also show in Fig. S7 D that the probability of a translated neuronal configuration under the receiving RBM is directly proportional to its probability under the original RBM. These results held across nearly all teacher–student pairs, with the exception of the outliers mentioned above (Fig. 4 E).

### Spatial structure is retained across fish

Finally, we asked whether the transferred activity patterns retained the spatial features of the source configuration. We computed the spatial correlation between the original and translated configurations at all time points. With the exception of the outliers described above (see Fig. S10), both *T* → *S* and *S* → *T* translations achieved significantly higher spatial correlations (median ≈ 0.55) than time-shuffled controls (median ≈ 0.3; Fig. 4 F–G), indicating that large-scale spatial motifs are preserved during transfer.

These results demonstrate that spontaneous neuronal activity patterns can be robustly translated across individuals through the shared RBM latent space. These translated configurations preserve key statistical and spatial features, are assigned high probability by the recipient model, and closely resemble the native activity observed in each fish. These findings indicate that the learned latent space encodes a shared and stereotyped repertoire of spontaneous brain-wide activity motifs. This framework offers a powerful new approach for comparing high-dimensional neuronal dynamics across animals, even in the absence of cell-by-cell correspondence.

## Discussion

In this study, we uncovered a shared latent structure underlying spontaneous brain-wide activity in larval zebrafish. By using Latent-aligned Restricted Boltzmann Machines (LaRBMs) across individuals, we identified a common latent space composed of functional assemblies: hidden units that represent recurrent co-activation patterns. The activity of one animal could be projected into this space and transferred into the neuronal space of another, preserving spatial structure and probabilistic consistency. These results reveal that spontaneous activity is structured by a shared repertoire of latent cell assemblies, suggesting conserved circuit-level constraints across brains.

Even in the absence of constraints that enforce a shared latent space, many co-activation motifs were spontaneously conserved across individuals. When RBMs were trained independently on individual fish, approximately 30% of the hidden units could be quantitatively matched across all animals based on spatial similarity. Visual inspection revealed that up to 50% displayed a conserved spatial density core, likely reflecting functional assemblies shaped by developmental or anatomical regularities. Although LaRBM does not uncover additional motifs beyond those of the teacher, it demonstrates that the teacher’s latent space acts as a shared non-linear functional basis, capable of representing spontaneous activity across diverse individuals. The success of this transfer supports the presence of stereotyped latent population codes that generalize beyond individual variability.

The RBMs’ shared hidden units exhibited spatial and functional consistency across individuals. This is remarkable because although student models were initialized using the spatial weight distribution of the teacher’s RBM, they were trained solely on functional activity without any spatial priors. Nevertheless, hidden units retained consistent spatial patterns and, in particular, pairwise correlations between hidden units were partially preserved across teacher and student. This correlation structure was not explicitly enforced during training, suggesting that the RBM uncovers latent activity motifs that recur across individuals. These motifs are functionally consistent, as reflected in preserved hidden-unit correlations, but also spatially conserved, despite the absence of explicit spatial constraints.

Moreover, the fact that the transferred activity patterns preserve low free energy in the recipient model indicates that the latent space captures not only structural regularities but also statistical plausibility, an essential property for generative models. Analogously to Helmholtz free energy in statistical physics, the free energy of an activity pattern measures the cost of maintaining a particular neural configuration (macrostate) by marginalizing over all compatible hidden configurations (microstates). Low free energy implies high probability under the model. This framework enables cross-individual comparison of brain-wide activity based on statistically grounded latent structure, allowing for quantitative evaluation of functional similarity grounded in principles from statistical physics.

Compared to other approaches for inter-subject alignment, such as anatomical registration, shared response modeling (SRM), hyperalignment, or contrastive autoencoders, our method offers several advantages. It is generative, allowing brain-wide activity patterns to be sampled, evaluated, and transferred probabilistically. It defines a probabilistic prior over spontaneous neural configurations, and after latent-alignment, this prior is shared across individuals, reflecting common structure in brain-wide activity. It is compositional, combining latent assemblies non-linearly to capture a richer repertoire of motifs than possible in linear subspaces. And it retains single-cell interpretability, unlike deep autoencoder methods. In addition, our approach is fully unsupervised and does not require stimuli, behavioral labels, or neuron correspondence. Lastly, RBMs are particularly well suited to our latent-alignment procedure because their hidden-unit prior *P*(*h*) is learned rather than fixed. It therefore adapts to the data and encodes shared co-activation motifs in latent space. By contrast, in methods such as variational autoencoders (VAEs), latent variables are typically regularized towards simple fixed priors, such as an isotropic Gaussian. Matching *P*(*h*) across individuals is therefore only a weak constraint in that setting, because similar latent priors do not guarantee alignment of the learned latent features themselves. A similar limitation applies to PCA, which makes latent directions comparable through linear rotation and scaling of covariance structure. However, because PCA standardizes only second-order statistics rather than learning shared latent structure, it does not ensure alignment of these latent features across individuals.

Although we focus here on spontaneous activity, the LaRBM framework is not intrinsically limited to it. One straightforward use case for stimulus-evoked recordings is to train LaRBMs on spontaneous segments of the recordings to learn an intrinsic dictionary of co-activation motifs, and then use the trained models to evaluate, reconstruct, and translate stimulus-evoked activity across animals. This provides a principled way to ask whether evoked responses recruit the same latent motifs as spontaneous dynamics, and whether the translated evoked patterns have comparable model free-energy/statistics in the target fish. More generally, training could also be performed on combined spontaneous-evoked datasets. However, stimulus-driven activity is likely to exhibit stronger temporal structure which would introduce higher-order moments between neuronal populations, overspecializing the model towards stimuli-evoked activity at the cost of spontaneous activity.

Despite these strengths, the current implementation of our approach has limitations. It assumes, on the mesoscale, a spatial organization of brain function. Although this assumption is appropriate for the larval zebrafish brains at 20–30 *µ*m resolution, it may not generalize to a more variable or less stereotyped system, such as the mammalian cortex. This is apparent in our analyses, where very few hidden units are assigned to the forebrain, particularly to the pallium, and activity in this region is not robustly translated between individuals. This might reflect a lack of consistent fish-to-fish stereotypy in forebrain spontaneous activity; however, it is more plausibly explained by weaker pairwise correlations within forebrain neuronal ensembles relative to those in the mid- or hindbrain. As a result, the model preferentially selects more strongly correlated units at the expense of pallial representations. A possible solution would be to train region-specific RBMs and recombine them into a single whole-brain model. Besides improving the representation of weakly correlated regions, this strategy would lower the memory required to train the larger models discussed below. In principle, the new LaRBM likelihood we introduce here should be sufficient to align latent spaces across individuals. Empirically, however, we have found that for whole-brain data the models do not converge with this new training paradigm alone, and thus require a *pre-training* phase, which we performed by spatially interpolating hidden units across individuals. With smaller, per-region RBMs, this pre-training might no longer be necessary.

Furthermore, the need to select a reference (teacher) fish introduces a source of bias. Although we observed robust generalization across multiple teacher choices, future extensions could mitigate this issue by implementing joint training or multi-teacher constraints. A symmetric, group-constrained approach, where all models are trained jointly with shared constraints, could enable the discovery of a richer and more balanced repertoire of conserved motifs. This would avoid teacher-specific bias and improve the statistical power to detect rare or subtle shared features. The resulting latent space could serve as a functional atlas onto which new data could be projected and compared across developmental stages, genotypes, or experimental paradigms. This framework could be used for functional phenotyping, where an individual’s deviation from the typical use of latent space may indicate neurodevelopmental or pathological alterations.

The emergence of spatially and statistically conserved latent motifs suggests that spontaneous dynamics are not arbitrary, but shaped by shared developmental programs and circuit architecture. The ability to identify and compare these motifs across individuals, without relying on anatomical correspondence or external stimuli, provides a new lens on how vertebrate brains are organized and how they differ. Together, our findings lay the foundations for a comparative neuroscience framework based on shared latent structure in spontaneous activity, enabling quantitative, model-based comparisons of brain-wide dynamics across individuals, developmental stages, or experimental conditions.

## Materials and Methods

### Data and Code Availability

All code, models, and post-processed data used in the present article are available from a centralized repository at *https://github.com/EmeEmu/ZebrafishSharedRBMs*.

Datasets and models are available for download at *https://doi.org/10.5281/zenodo.19066472*. Raw data is available upon request (approximately 400GB).

Calcium imaging pre-processing was performed using previously published protocols and software implemented in MATLAB (Mathworks) [31, 43, 26, 38].

Morphological Registrations was performed using the Advanced Normalization Tools (ANTs) [39].

The Restricted Boltzmann Machine modeling is based on the general-purpose Julia package *https://github.com/cossio/RestrictedBoltzmannMachines.jl* [11]. An extension to this package implementing the LaRBM procedure is available at *https://github.com/BiRBMs2024/BiTrainedRBMs.jl/tree/main*.

### Fish Husbandry

All experiments were performed on nacre Tg(elavl3:H2B-GCaMP6f) Danio *rerio* larvae aged 5 to 6 days post-fertilization (dpf). Larvae were reared in groups of 20-30, in Petri dishes in embryo medium (E3) on a 14/10 hour light/dark cycle at 28^◦^*C*, and were fed 1*mL* of rotifers at 10^6^*/ℓ* daily from 5 dpf.

Larvae were screened at 4 and 5 dpf for GCaMP6f expression, checking for fluorescence intensity and uniform expression patterns. At 5 dpf, larvae were also screened for correct inflation of the swim bladder.

The experimental protocols were approved by Le Comitée d’Ethique pour l’Expérimentation Animale Charles Darwin C2EA-05 (02601.01 and #32423-202107121527185 v3).

### Experimental Protocol

#### Fish Preparation

Larvae were paralyzed by immersion in a 1*mg/mL* solution of alpha-bungarotoxin (Invitrogen™ B1601) for one to two minutes in groups of 5 fish. After treatment, the larvae were washed and placed in E3 solution for 20 minutes. Startle response and blood flow were then checked to assess correct paralysis.

A single larvae is then mounted tail first in a capillary (Wiretrol® II 25 − 50*µL*) in 2% low melting point agarose (Invitrogen™ 16520-100), and placed in the imaging tank, in E3 solution, under the microscope for an habituation period of 1 hour.

#### Imaging

Spontaneous neural activity was recorded using Light-Sheet Microscopy, at 2.5 volumes per second during 25 minutes, with 25 z-planes separated by 10*µm* [31].

A total of 17 fish were recorded for this study. Of these, 3 fish were discarded due to failing paralysis, 2 died during the recording, and 4 presented uncorrectable drift in z; leading to 9 datasets which could not be exploited. Of the 8 remaining fish, another 2 fish were discarded due to abnormally low neuronal activity, which we suspect could have been caused by an excessive paralysis. This leaves 6 datasets which were exploitable and presented *normal* brain activity. No further selection was performed, and all 6 fish were pre-processed and used in the present study.

Fish 1 and 3 were randomly chosen as examples in all main figures, with Fish 3 as teacher and Fish 1 as student in all relevant figures (see Tab. 1).

#### Data pre-processing

Image pre-processing was performed offline using MATLAB according to previously published protocols [31, 43, 26].

Two dimensional drift correction was applied to each imaging layer to correct for movements of the sample during recording. Watershed was then used for cell segmentation, and a mean fluorescence value was computed from each cell ROI *i* and frame *t* : *F*_*it*_. This was then normalized to Δ*F/F*_*it*_ = (*F*_*i*_ − ⟨*F*_*i*_⟩_*t*_)*/*(⟨*F*_*i*_⟩_*t*_ − *F*_0_), where ⟨*F*⟩_*t*_ is the baseline fluorescence of neuron *i* and *F*_0_ is the image-wide background fluorescence.

The Δ*F/F*_*it*_ was then deconvolved into binarized spike trains *s*_*it*_ ∈ [0, 1] using Blind Sparse Deconvolution (BSD) [38], using the calcium kernel time constants for GCaMP6f previously inferred using BSD [26, 41] : rise time 0.15s and decay time 1.6s. This method was used over other published methods as it not only deconvolves the fluorescence into spike trains, but also provides a rationale for the binarization of this train.

The first 4 minutes (600 frames) of each recording were excluded. The fish were accustomed to the imaging chamber for 1 hour with the laser turned off to avoid photobleaching. When the laser is turned on at the start of recording, the mean calcium signal rises sharply and then decays to a stable level within about 1 min. By discarding the initial 4 min, we ensure that all analyzed data come from the steady-state phase of neuronal activity under continuous illumination.

Samples of the binarized whole-brain activity for each fish can be seen in Movie S1.

#### Morphological Registration

For each recording, a high-resolution stack was acquired with 250 z planes separated by 1*µm* and a camera exposure time of 200*ms* per layer. A low-resolution stack was then created by averaging all frames of the functional recording, and then rigidly registered to the high-resolution stack using the Advanced Normalization Tools (ANTs) [39].

The high-resolution stacks of all fish were then registered together using both affine and warp transformations to create a single reference space in which all segmented neurons could be projected. The ANTs command and parameters were adapted from Vladimirov et al. [42], and are given bellow.

~~~
antsRegistration
     --float 1 --dimensionality 3 --interpolation WelchWindowedSinc --use-histogram-matching 0
     --output [outputdir/LowRes_TO_HighRes_,outputdir/LowRes_TO_HighRes.nrrd]
     --initial-moving-transform [HighRes.nrrd,LowRes.nrrd,1]
     --transform Rigid[0.1] --metric MI[HighRes.nrrd,LowRes.nrrd,1,32,Regular,0.25]
     --convergence [200x200x200x0,1e-8,10] --shrink-factors 12x8x4x2 --smoothing-sigmas 4x3x2x1vox
     --transform Affine[0.1] --metric MI[HighRes.nrrd,LowRes.nrrd,1,32,Regular,0.25]
     --convergence [200x200x200x0,1e-8,10] --shrink-factors 12x8x4x2 --smoothing-sigmas 4x3x2x1vox
antsRegistration
     --float 1 --dimensionality 3 --interpolation WelchWindowedSinc --use-histogram-matching 0
     --output [HighRes_TO_Reference_,HighRes_TO_Reference.nrrd]
     --initial-moving-transform [Reference.nrrd,HighRes.nrrd,1]
     --transform Rigid[0.1] --metric MI[Reference.nrrd,HighRes.nrrd,1,32,Regular,0.25]
     --convergence [200x200x200x0,1e-8,10] --shrink-factors 12x8x4x2 --smoothing-sigmas 4x3x2x1vox
     --transform Affine[0.1] --metric MI[Reference.nrrd,HighRes.nrrd,1,32,Regular,0.25]
     --convergence [200x200x200x0,1e-8,10] --shrink-factors 12x8x4x2 --smoothing-sigmas 4x3x2x1vox
     --transform SyN[0.05,6,0.5] --metric CC[Reference.nrrd,HighRes.nrrd,1,2]
     --convergence [200x200x200x200x10,1e-7,10] --shrink-factors 12x8x4x2x1
     --smoothing-sigmas 4x3x2x1x0vox
antsApplyTransforms
     --dimensionality 3 --float --input LowRes.nrrd --reference-image Reference.nrrd
     --output outputdir/LowRes_TO_Reference.nrrd --interpolation WelchWindowedSinc --default-value 0
~~~

### Restricted Boltzmann Machines

#### Definition

Restricted Boltzmann machines (RBM) [18] are two-layer energy-based models, over *N* visible units **v** = (*v*_1_, …, *v*_*N*_ ) representing the neural activity with binary values *v*_*i*_ ∈ {0, 1}, and *M* real-valued hidden units **h** = (*h*_1_, …, *h*_*M*_ ).

The RBM defines a probability distribution over configurations of all the units given by:

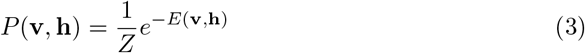

where *Z* is a normalization constant known as the partition function, and *E*(**v, h**), the energy, is given by:

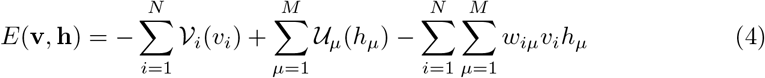

The functions 𝒱_*i*_(*v*_*i*_) and 𝒰_*µ*_(*h*_*µ*_) are potentials biasing the activities of single units, while the weights *w*_*iµ*_ account for interactions between visible and hidden units. In the case of neuronal data, we take 𝒱_*i*_(*v*_*i*_) = *g*_*i*_*v*_*i*_ with *g*_*i*_ called the field. Following van der Plas et al. [41], we use dReLU potentials for the hidden units,

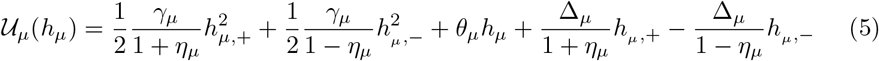

with real parameters *γ*_*µ*_, *η*_*µ*_, Δ_*µ*_, *θ*_*µ*_, satisfying *γ*_*µ*_ *>* 0. Here *h*_*µ*,+_ = max(*h*_*µ*_, 0) and *h*_*µ*,−_ = min(*h*_*µ*_, 0).

To obtain the probability of neural activity, the hidden units are marginalized. In this way, one defines a marginal likelihood over the visible units:

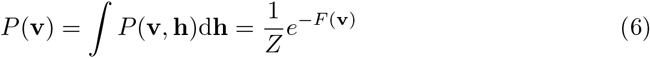

where *F*(**v**) is a free energy function that incorporates the effective interactions induced by the marginalized hidden variables,

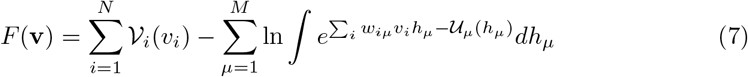

#### Standardized RBMs

Following [12], we employ a generalized *centering trick* [28], where the activity of the hidden units is not only centered, but also standardized.

The Standardized Restricted Boltzmann machine (StdRBM) is a re-parameterization of the RBM, where the energy function is defined as follows:

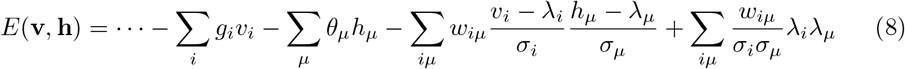

where we only show the fields for the unit potentials for simplicity. During training, the parameters *λ*_*i*_, *λ*_*µ*_ and *σ*_*i*_, *σ*_*µ*_ track a moving average of the mean and standard deviations of the corresponding unit activities. After training, the StdRBM can be converted into an equivalent RBM by the transformations:

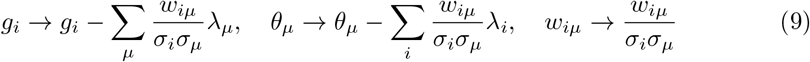

As has been observed empirically in previous works [28, 12, 25], such centering and standardization of unit activities leads to improved training convergence and stability.

#### Training

All RBM parameters (*g*_*i*_, *γ, η*, Δ, *θ, w*_*iµ*_) are trained to maximize the likelihood of a neural activity dataset. If a data set of neural activity recordings is given as 𝒟 = {**v**^*t*^}, where *t* is the index of the sample (or time), then the RBM is trained by maximizing:

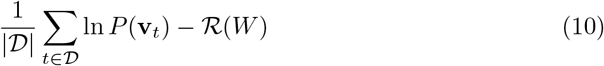

Here, ℛ (*W* ) is a regularization term applied over the RBM weights. Following van der Plas et al. [41], we employ an *L*_1_ regularization for single-cell RBMs,

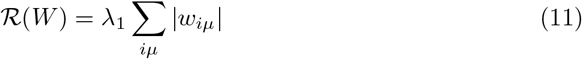

where *λ*_1_ controls the regularization strength. As in van der Plas et al. [41], we set *λ*_1_ = 0.02, and *M* = 100 hidden units. The optimization is carried out by gradient ascent of Eq. (10). We use the ADAM algorithm [22].

#### Sampling and Reconstruction

Once trained, the RBM can generate synthetic neuronal configurations by alternating between sampling hidden states from *P*(*h* | *v*) and visible states from *P*(*v* | *h*), forming a Gibbs sampling chain. This process allows the model to reconstruct expected activity patterns from observed configurations and explore the distribution it has learned. Movie S2 illustrates a single Gibbs sampling step and the resulting reconstruction ⟨*v*⟩ from an observed configuration in Fish 1.

#### Evaluation

Trained RBMs are evaluated on their ability to reproduce data statistics. As they are trained to maximize the log-likelihood of the data, a well trained RBM is expected to generate data with statistics ⟨*v*_*i*_⟩, ⟨*h*_*µ*_⟩, and ⟨*v*_*i*_*h*_*µ*_⟩ matching the corresponding empirical data statistics. We also evaluate the RBMs’ ability to reproduce the second-order statistics ⟨*v*_*i*_*v*_*j*_⟩ − ⟨*v*_*i*_⟩ ⟨*v*_*j*_⟩ and ⟨*h*_*µ*_*h*_*ν*_⟩ − ⟨*h*_*µ*_⟩ ⟨*h*_*ν*_⟩ to ensure that first order interactions between neurons are captured by the model [41]. An RBM has to reproduce well all five statistics to be considered usable in our use case.

Measurement of statistics matching between model-generated and empirical data is done using the normalized Root Mean Squared Error (nRMSE) as introduced by van der Plas et al. [41] (see Measuring identity). This measurement is standardized such that a value of 1 corresponds to shuffled data statistics, and a value of 0 corresponds to the optimal matching expected from the difference between train and test sets. The evaluation of an RBM is thus represented by a vector of length 5 containing the nRMSE for all five statistics, and multiple RBMs can be compared using the *L*_∞_ norm.

#### Inferred neuronal couplings

van der Plas et al. [41] showed that an effective coupling matrix *J*_*ij*_ could be inferred from a trained RBM model. This is done by manually perturbing the activity of each neuron and quantifying the impact on other neurons by evaluating the marginal distribution *P*(**v**). Given a neuronal configuration **v**, the coupling is then defined as

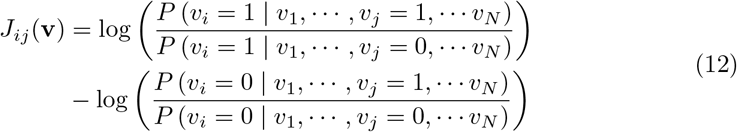

The overall coupling matrix is then obtained from all empirical neuronal configurations as :

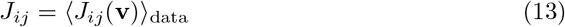

However, calculating *J*_*ij*_(**v**) is computationally demanding, and we use the following approximation introduced by van der Plas et al. [41]:

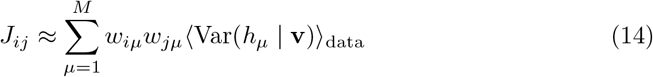

#### Graphical ordering of hidden units

Hidden units have no intrinsic order in an RBM (the model is invariant to permutation of hidden-unit indices). For visualization purposes only, we therefore reordered hidden units in figures by hierarchical clustering of their pairwise activity correlation matrix corr(*h*_*µ*_(*t*), *h*_*ν*_ (*t*)), computed from the inferred hidden-unit time series **h**(*t*) = 𝔼_**h**|**v**_[**v**(*t*)].

We then applied optimal leaf ordering to the resulting dendrogram to obtain a one-dimensional ordering that places highly correlated hidden units adjacent. This reordering is only used to improve figure readability and does not affect training, inference, or any quantitative analyses.

### Voxelised RBMs

#### Voxelisation

To obtain a coarse-grained description of the datasets, we voxelize the brain into cubes of side *v* (typically *v* = 20*µm*). This grid is defined in the common registered brain space. Neurons are assigned to the voxels after segmentation, according to their center of mass. The activity of each voxel for each fish is then defined as :

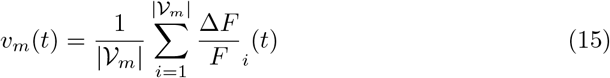

where *v*_*m*_ is the activity of voxel *m* containing the set of neurons *V*_*m*_ and 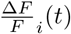 is the normalized fluorescence signal of neuron *i* at time *t*.

This activity is then normalized for each fish *f* and each voxel using z-score :

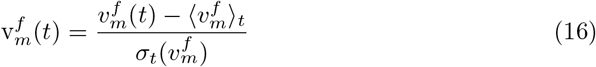

This process yields a series of voxelized activities 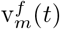 where each voxel *m* corresponds to the same anatomical position across all fish.

To ensure that the z-scored activity of each voxel is well defined, we discard voxels for which 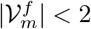 for at least one fish.

#### Training

Contrary to the neuronal RBMs described above (Restricted Boltzmann Machines), in the case of voxelized data, we used Standardized RBMs with gaussian visible units, as the voxelized activity approximately follows a normal distribution (see Fig. S3 C).

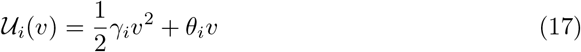

We found empirically that for voxelized RBMs, *L*_1_ regularization produced weight matrices where the majority of hidden units were decoupled from the visible layer.

Hence we use the 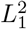 regularization introduced by Tubiana, Cocco, and Monasson [37] :

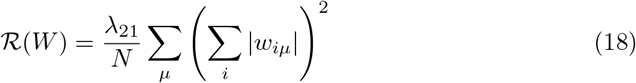

where *N* is the number of voxels and *λ*_21_ is the regularization factor.

We performed a cross-validation over the number of hidden units *M* and regularization factor *λ*_21_ on Fish 1 (Fig. S3 A). We identified 3 sets of hyperparameters which we then tested on the concatenated voxelized data of all 6 fish in the dataset (Fig. S3 B), and chose *M* = 40 and *λ*_21_ = 0.1.

The rest of the training parameters are listed here :

- 20000 gradient updates.
- 50 Markov Chain steps between gradient updates.
- 100 time points per batch.
- 5· 10^−4^ learning rate for the first 1*/*4 of the training. The learning rate is then annealed with a geometrical decay to reach 10^−5^ at the end of training.

### Latent-aligned Restricted Boltzmann machines (LaRBMs)

#### Teacher RBMs

Teacher standardized RBMs were constructed and trained from whole-brain binarized neuronal activity with the following parameters :

- *M* = 100 hidden units (instead of either 200 or 100 in van der Plas et al. [41] as most of our fish have weak activity in the optic tectum).
- *λ*_1_ = 0.02 for weight L1 regularization (as obtained by van der Plas et al. [41] after cross-validation).
- 200000 gradient updates.
- 15 Markov Chain steps between gradient updates.
- 256 time points per batch.
- 5 · 10^−4^ learning rate for the first 1*/*4 of the training. The learning rate is then annealed with a geometrical decay to reach 10^−5^ at the end of training.

See Supplementary Hyper-parameters selection for a more detailed discussion of the choice of hyperparameters and the impact on the main results presented here.

Before training, individual fish datasets were split into a 70% training and 30% validation sets using the method described by van der Plas et al. [41]. The validation set was used to evaluate RBM-generated vs empirical data statistics as described in Evaluation.

For every fish, we trained 10 RBMs and kept the one which performed best as described in Evaluation (see Fig. S5 for all training statistics). Indeed, we found that convergence can be very variable, with some models converging satisfactorily, and other either converging poorly or not converging at all.

#### Student LaRBM Initialization

Student LaRBMs are initialized from the teacher with the following steps.

First, the fields *g*_*j*_ of the visible layer are estimated from logit of the average neuron activity 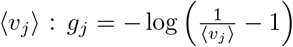. The visible scaling (*σ*_*j*_) and bias (*λ*_*j*_) parameters used to standardize the RBM are also estimated from neuronal activity as :

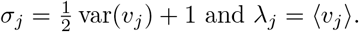

Second, hidden potentials 𝒰_*µ*_(*h*_*µ*_) (i.e. all parameters of the dReLU *γ*_*µ*_, *η*_*µ*_, Δ_*µ*_, *θ*_*µ*_) are copied directly from the teacher RBM.

Third, the teacher weight matrix *w*_*iµ*_ is spatially interpolated at the locations of student neurons. To do so, we first build a 3D map of weights for each hidden unit *µ* :

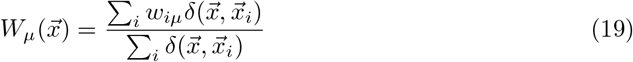

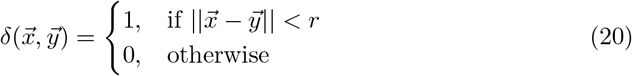

where 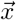 is any 3D location in shared brain space, 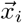 is the 3D location of teacher neuron *i*, and 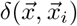 is a boolean ball of radius *r* = 4*µm*, the average size of neurons.

These maps are then convolved with a gaussian kernel 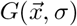 of width *σ* = 4*µm* to smooth out local discontinuities : 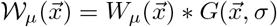. Because weight matrices are very sparse, this smoothing step has a tendency to attenuate the weight map. We therefore add a bias :

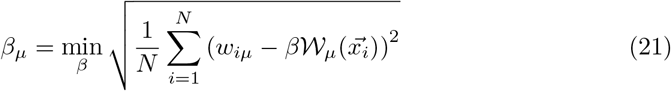

Finally, a weight matrix for the student can be created by evaluating 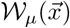 at the locations 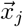 of neurons from the student fish :

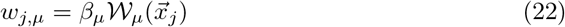

### Student training

Once initialized, student RBMs are trained with the same set of parameters as the teacher, except for the number of gradient updates for which we found 1/10th (20000) to be sufficient for convergence.

In absence of any constraints, the space of representations learned by the student can drift arbitrarily far from the one of the teacher. This is undesirable because we cannot compare the hidden unit activities from the teacher and the student. In order to avoid this, we introduce a new training paradigm to maintain an alignment between the hidden units of the student and the teacher. Mathematically, we would like the marginal distribution of hidden unit activities, *P*(**h**) = ∑_**v**_ *P*(**v, h**), to be similar in the teacher and the student. To achieve this, we first sample hidden unit activity from the teacher RBM to create a dataset of hidden unit activities, 𝒯 = {**h**_*t*_} from the teacher. The student is trained following the modified objective:

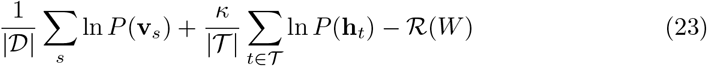

The first and last terms are the same appearing in the original training objective, Eq. (10). The second term is the average log-likelihood of the teacher hidden unit activity evaluated under the student. By maximizing this term, we force the student to align its marginal hidden unit distribution to match that of the teacher. The coefficient *κ* controls the relative importance of this term: for *κ* = 0, we recover the original training objective of Eq. (10), while for *κ* large, the student RBM is forced to match the teacher hidden unit statistics more closely.

### Translating neuronal activity patterns across individuals

In principle, translating activity from the visible layer of one RBM, through the hidden layer, to the visible layer of another RBM should be done via sampling the hidden layer given the first fish visible layer, and for each sample, sampling the second fish visible layer given the hidden layer, and then averaging over all samples, which yields 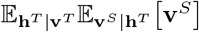. However, in practice, this is computationally intensive and thus impractical in our use case. Instead, we use 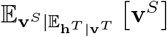 as we have found empirically that for a high number (∼ 1000) of samples the two method give similar results (see Fig. S8 A-D).

This similarity can be explained with a Taylor expansion on 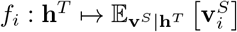

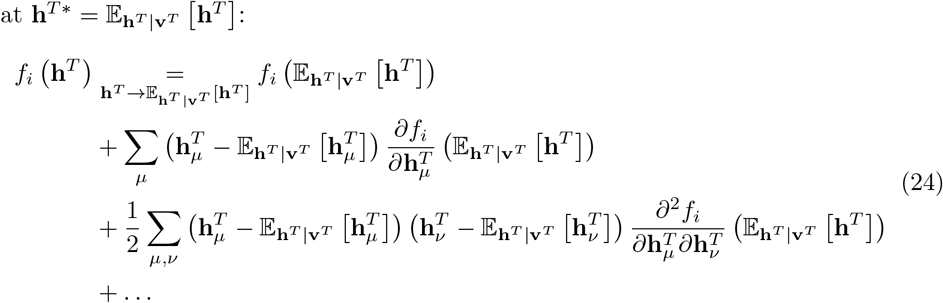

Because of the conditional independence of **h**^*T*^ given **v**^*T*^,

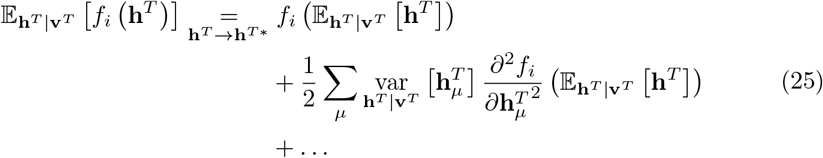

Since 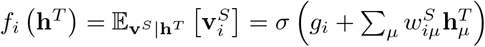, where 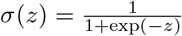, we find

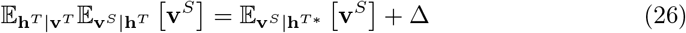

where

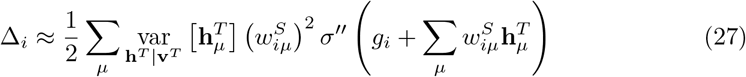

We show on Fig. S8 that Δ is indeed negligible compared to 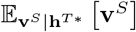

### Measuring identity

As introduced by van der Plas et al. [41], we quantify the goodness of fit to the identity using the normalized Root Mean Squared Error (nRMSE). It is defined as :

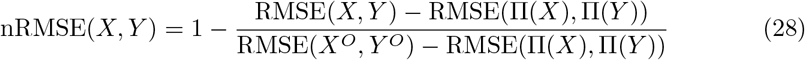

where *X* and *Y* are two vectors of same length, RMSE 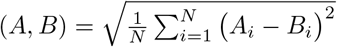 the standard Root Mean Squared Error, (*X*^*O*^, *Y* ^*O*^) is the pair of optimal vectors corresponding to the best expected fit of *X* and *Y*, and Π is a shuffling operator such that Π(*A*)_*i*_ = *A*_*π*(*i*)_.

Unlike the standard RMSE, this measure is independent of the length of the vectors *X* and *Y*, and is easily interpretable. Indeed, a value of nRMSE(*X, Y* ) = 1 means that *X* and *Y* are as uncorrelated as shuffled vectors, while nRMSE(*X, Y* ) = 0 means that *X* and *Y* are as identical as can be expected from the optimal (*X*^*O*^, *Y* ^*O*^).

In the case where no optimal (*X*^*O*^, *Y* ^*O*^) can be defined other than *X*^*O*^ = *Y* ^*O*^, RMSE(*X*^*O*^, *Y* ^*O*^) = 0 and therefore :

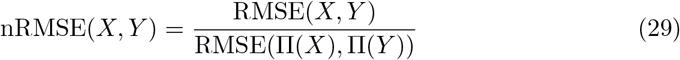

nRMSE(*X, Y* ) = 0 is then interpreted as *X*_*i*_ = *Y*_*i*_ ∀*i*.

### Measuring Spatial Similarity

To compare how a spatially defined variable (such as neuron weight or firing rate) is distributed across two different fish, we define a spatial similarity measure between animals A and B.

Let *ϕ*^*A*^ and *ϕ*^*B*^ be two spatial fields that assign scalar values 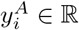 and 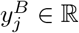 to neurons at 3D positions 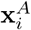 and 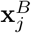 in fish A and B, respectively: 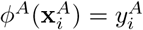 and 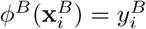. We wish to estimate how well *ϕ*^*A*^ aligns with *ϕ*^*B*^.

Since neurons are not spatially matched between individuals, we interpolate the values 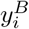 from fish B to the locations of neurons in fish A using Gaussian kernel smoothing :

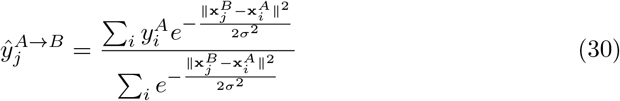

where 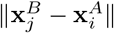 is the euclidean distance between neuron *i* from fish A and neuron *j* from fish B, and *σ* is the standard width of a gaussian kernel, which we take to be a neuron size 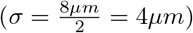. This provides an estimate of what the spatial distribution of *y* from fish A would look like in fish B.

We can then evaluate the similarity between the two spatial fields *ϕ*^*A*^ and *ϕ*^*B*^ by computing the Pearson correlation coefficient between 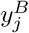 and 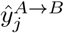 :

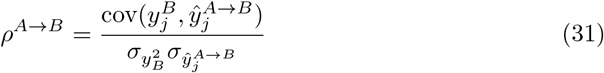

In practice, the Person correlation is used over other identity tests such as the nRMSE (see Methods Measuring identity) because our observables tend to be sparse and the gaussian interpolation will thus naturally lower *ŷ* (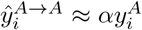 with α < 1).

In the case of the weight maps presented in Fig. 3 B-F, *ρ*^*A*→*B*^ is only computed for neurons whose weights are significant (*>* 10^−5^), as we are only interested in whether student maps are compatible with teachers, and the regularization imposed on weights during RBM training will force a non-spatial linearity in weight distribution.

### Measuring Diagonal Dominance

In Fig. 3 C-D, we ask whether the spatial distribution of hidden units’ weights is maintained between teacher and student RBMs, in a way that allows for their unambiguous pairwise identification. In other words, we ask whether the spatial distribution of 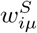 is closer to 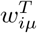 than to any other hidden unit 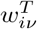with ν≠ µ.

To do so we first compute the matrix 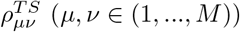 of spatial correlation (see Methods Measuring Spatial Similarity) between the weight map of hidden unit *µ* in the teacher RBM, and hidden unit *ν* in the student RBM.

Our hypothesis is confirmed not only if each teacher hidden unit *µ* is most similar to its corresponding student unit (i.e., if arg max_*ν*_ 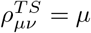), but also if this match is significantly better than any other (i.e., 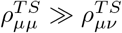 for all *ν≠* µ).

To test for this systematically, we perform a bootstrap analysis. We start by computing a *distance* between the diagonal and the next largest element of the matrix :

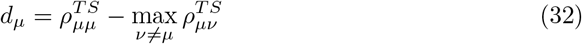

We next compute the same distance for *n* = 5000 random row permutations of the matrix 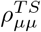:

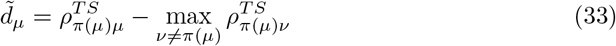

where *π*(*µ*) is a random permutation of 1, …, *M* (see Fig. S7 C for an example). We can then compute a p-value measuring the probability that 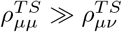 could be explained by random permutations of the matrix :

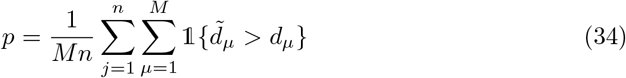

It is to be noted that, as some hidden units are completely disconnected from the visible layer (*w*_*iµ*_ < 10^−5^, ∀ *i*), some weight maps are undefined. These are therefore omitted from this measurement.

## Acknowledgments

We would like to thank Simona Cocco, Rémi Monasson, and Jérôme Tubiana for helpful discussions and insights. As well as Ans Imran for his help during his internship.

We would also like to thank Abdelkrim Mannioui, Marco Amaral, Edouard Manzoni and the rest of the team at the fish facility for taking care of the animals, as well as Malika Pierrat and Laura Bacerra-Zapata for taking care of the administration and contracts.

This project has received funding from CNRS, Sorbonne University, and the European Research Council (ERC) under the European Union’s Horizon 2020 research innovation program grant agreement number 715980. M.DK had a PhD fellowship from the École Doctorale Frontière de l’Innovation en Recherche et Education - Programme Bettencourt. J.FdCD was supported by a CNRS Chaire de Professeur Junior (CPJ) starting package ANR-24-CPJ1-0172-01. G.FB had a PhD fellowship of the i-Bio Initiative from the Idex Sorbonne University Alliance.

## Supporting information

### Supporting movies

**Movie S1:** Spontaneous whole-brain activity (six fish). Spontaneous, whole-brain, single-neuron binarized spiking activity from six larval zebrafish (Fish 1–Fish 6) recorded with Light-Sheet Microscopy. Shown as a dorsoventral projection, playback is 4× real time.

**Movie S2:** RBM sampling and reconstruction (Fish 1). Sampling of a Restricted Boltzmann machine (RBM) trained on the binarized, whole-brain activity of Fish 1. Left: observed neuronal configuration v. Middle: one RBM Gibbs sampling step *v* →*h* →*v*. Right: RBM reconstruction ⟨v⟩ expected from the observed configuration. Shown in dorsoventral projection.

**Movie S3:** Teacher→Student translation (Fish 3 →Fish 1). Translation of neuronal configurations from the teacher RBM (Fish 3) to the student RBM (Fish 1) through the shared latent space. Left: observed configuration v (teacher, Fish 3). Middle: teacher reconstruction. Right: configuration translated into the student’s neuronal space. Dorsoventral projection, playback 4× real time.

**Movie S4:** Student→Teacher translation (Fish 1 →Fish 3). Same as Movie S3, but translating from student to teacher.

All movies are shown in dorsoventral projection. Movies S1, S3, and S4 are accelerated 4×.

## Supporting figures

**Fig S1.**
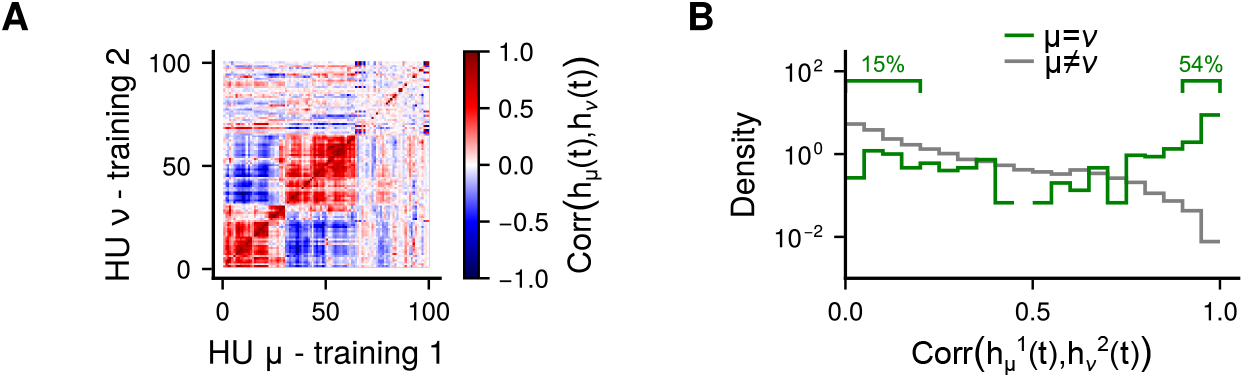
Supplementary for Fig. 1. **A**: Pairwise Pearson Correlation between hidden activity *h*_*µ*_(*t*) = 𝔼 [**h**| **v**_*t*_] and *h*_*ν*_ (*t*) of two example RBMs trained on the same neuronal recording. The matrix rows have been reordered to obtain the best alignment between the two trainings. **B**: Distribution of correlations from panel A, for best alignments pairs (*ν* = µ, green) and other pairs (ν≠ µ, grey)

**Fig S2.**
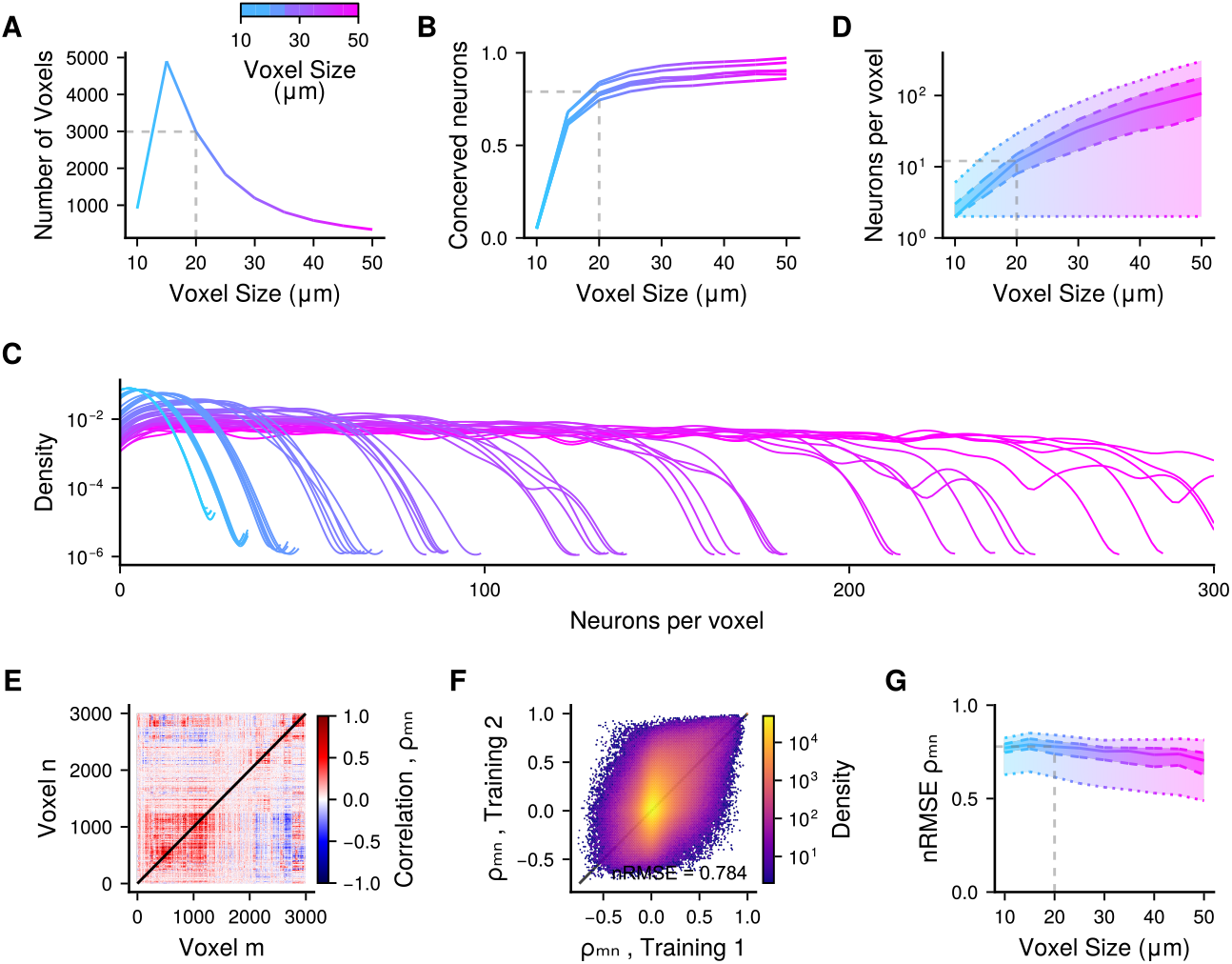
Supplementary for Fig. 2 investigating the impact of voxel size. **A**: Number of populated voxels (each voxel needs to contain ≥2 neurons for each fish, see Methods Voxelised RBMs) as function of voxel size. **B**: Fraction of neurons conserved after voxelization. Each line is a fish. **C**: Distributions of the number of neurons per voxel, for each voxel-size tested (same colors as in panel A), and for each fish (one line per fish). **D**: Median (solid line), 25-75% range (dashed lines), and min-max range (dotted lines) of the number of neurons per voxel (all fish combined). **E**: Pairwise Person correlation coefficients ρ_*mn*_ between voxels *m* and *n* for an example fish at 20*µm* voxel size. **F**: Stereotypy in ρ_*mn*_ between 2 example fish at 20*µm* voxel size. **G**: nRMSE (see Methods Measuring identity) of ρ_*mn*_ between 2 example fish as a function of voxel size.

**Fig S3.**
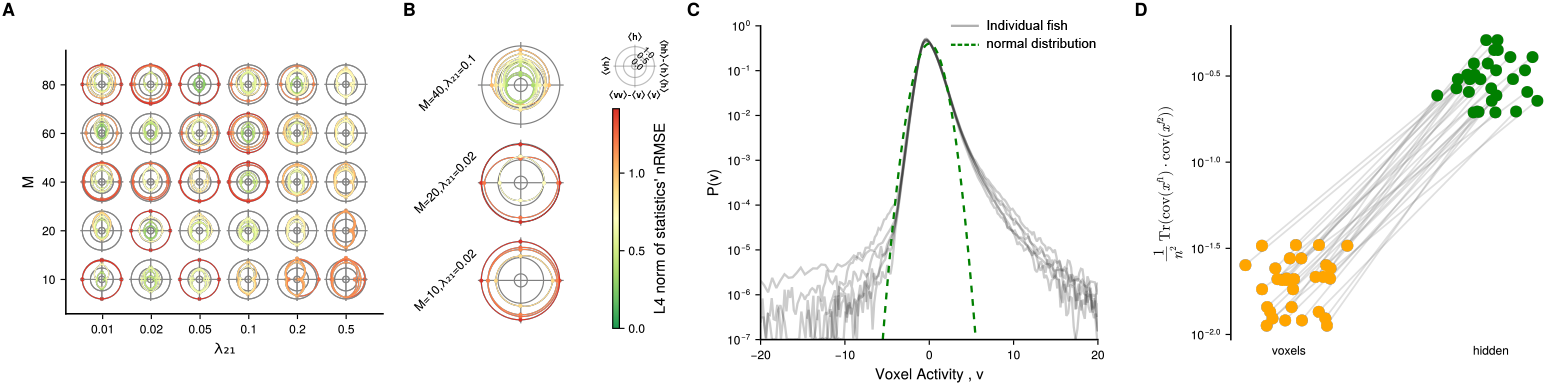
Supplementary for Fig. 2 cross-validation of voxelized RBM hyperparameters. **A**: nRMSE (see Methods Measuring identity) of all four moments (see Methods Evaluation) used to evaluate RBM convergence ( ⟨v⟩ is omitted as voxelized data is normalized by z-score). Each RBM is represented by a line in polar coordinates where every ray corresponds to a moment *m* ∈ {⟨**vh**⟩, ⟨**h**⟩, ⟨**vv**⟩ − ⟨**v**⟩⟨**v**⟩, ⟨**hh**⟩ − ⟨**h**⟩⟨**h**⟩}, and the ray length corresponds to the nRMSE_*m*_(data, generated) between empirical data and data generated by the RBM. Five training were performed and evaluated per hyperparameter pair (*M, λ*_21_), where *M* is the number of hidden units, and λ_21_ is the regularization factor (see Methods Training). All trainings were done on voxelized data from Fish 1. **B**: Same panel A for 3 sets of hyperparameters, and for the concatenated voxelized data of all 6 fish in the dataset. **C**: Distributions of voxelized activity values, with one grey line per fish. Normal distribution in green (dashed). **D**: Fish to fish alignment of eigen vectors in voxelized and hidden space : 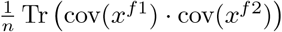, where *x*^*f1*^ is activity of one fish, either the voxel activity *v*_*tn*_ or the hidden units activity *h*_*tn*_, *f* 1 and *f* 2 are two different fish, and *n* is either the number of voxels or the number of hidden units.

**Fig S4.**
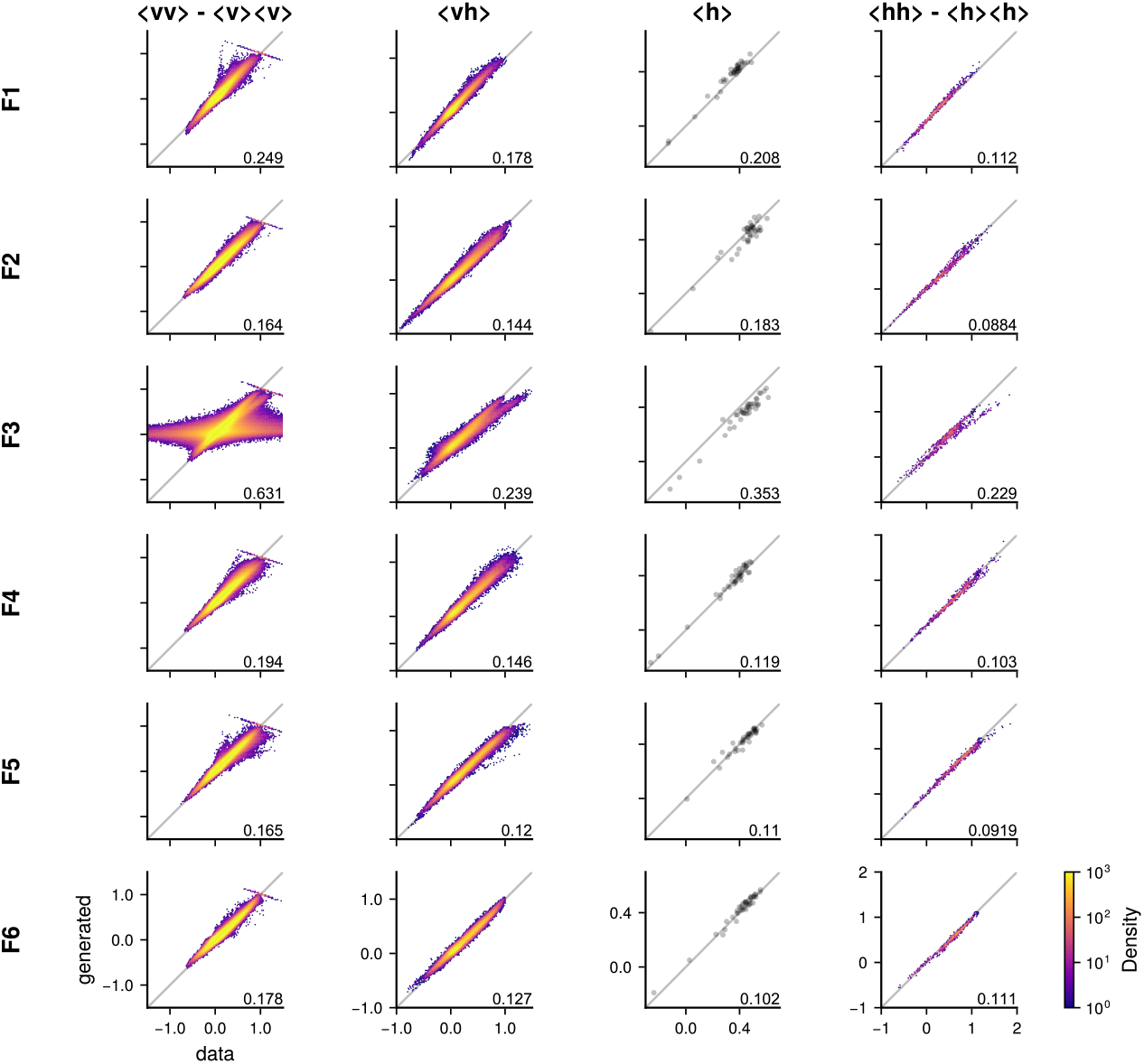
Supplementary for Fig. 2 showing the voxelized training statistics for each fish. Identity plots for all 4 moments (see Methods Evaluation) used to evaluate voxelized RBM convergence (notice that moment ⟨v⟩ is not represented as voxelized activity is normalized by z-score, see Methods Voxelised RBMs) for each fish.

**Fig S5.**
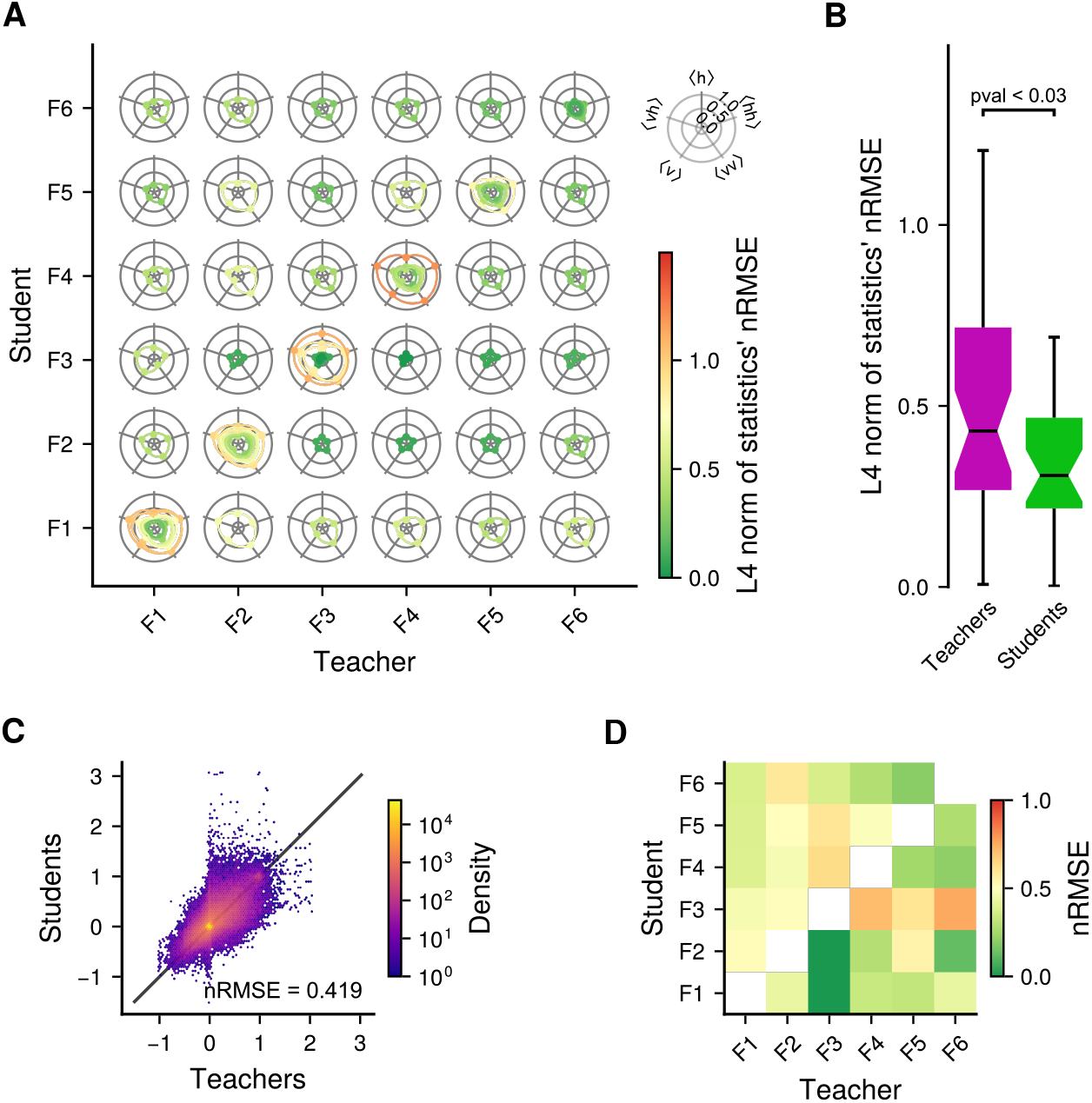
Supplementary for Fig. 3 showing the training statistics of every teacher-student pairs. **A**: nRMSE (see Methods Measuring identity) of all five moments (see Methods Evaluation) used to evaluate RBM convergence. Each RBM is represented by a line in polar coordinates where every ray corresponds to a moment *m* ∈ {⟨**v**⟩, ⟨**vh**⟩, ⟨**h**⟩, ⟨**vv**⟩ − ⟨**v**⟩⟨**v**⟩, ⟨**hh**⟩ − ⟨**h**⟩⟨**h**⟩}, and the ray length corresponds to the nRMSE_*m*_(data, generated) between empirical data and data generated by the RBM. The line color corresponds to the infinity norm of all five statistics. RBMs along the diagonal correspond to teacher RBMs trained classically (see Methods Standardized RBMs, 10 repetitions). Of-diagonal RBMs are students. (column 3 of this graph corresponds to Fig. S6) **B**: Comparison of training accuracy between teachers (left) and students right, measured as the L4 norm of the moments nRMSE in panel A. The p-value was computed with a one-tailed Mann-Whitney U test. **C**: Identity plot between teacher and student hidden unit pairwise covariance ⟨**hh**^⊤^⟩ − ⟨**h**⟩⟨**h**^⊤^⟩, for all teacher-student pair combined. **D**: nRMSE of ⟨**hh**⟩ − ⟨**h**⟩⟨**h**⟩ for every teacher-student pair.

**Fig S6.**
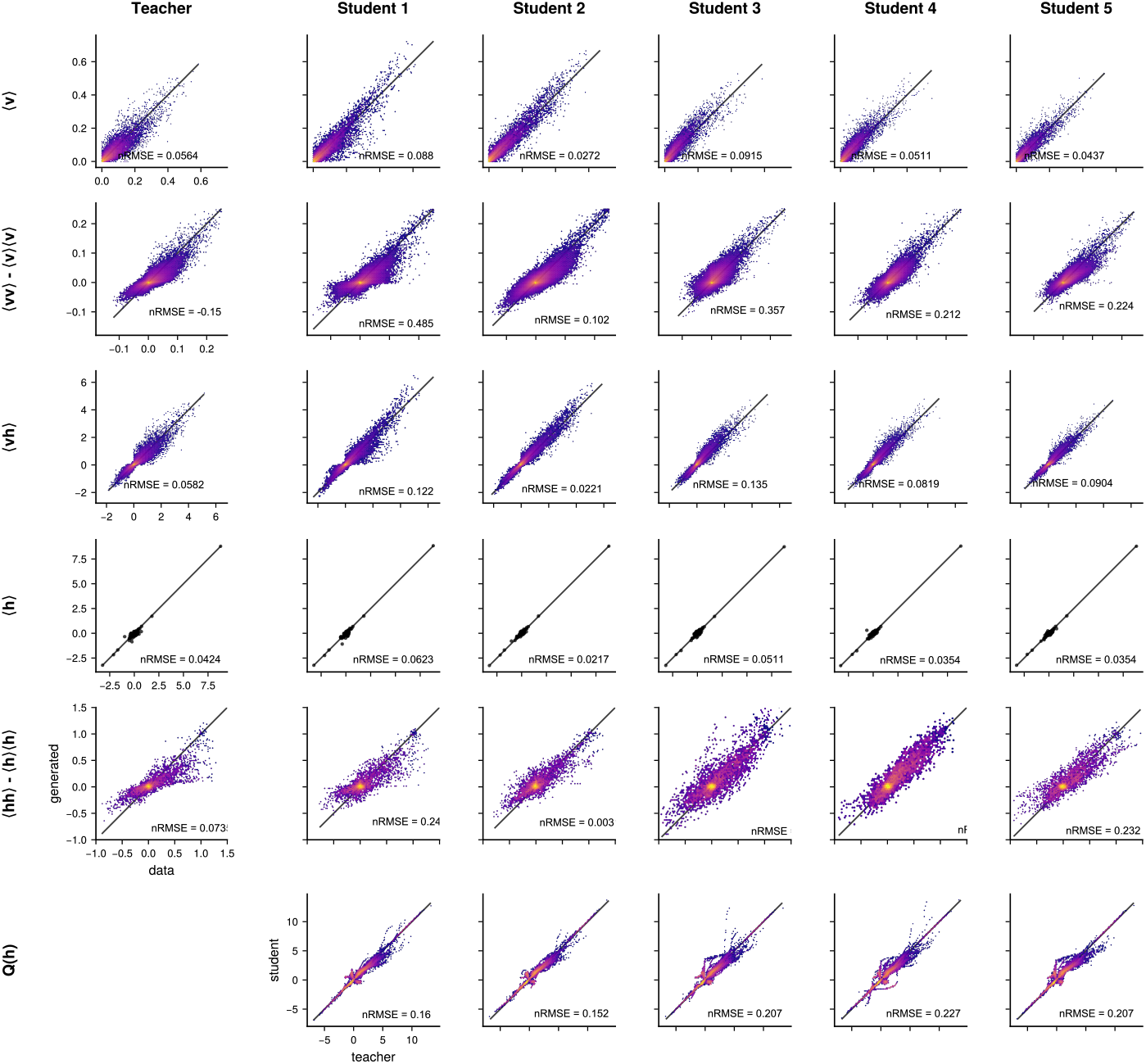
Supplementary for Fig. 3 showing the training statistics and hidden distribution of all students of the example teacher fish. Identity plots for all five moments (see Methods Evaluation) used to evaluate RBM convergence and quantile-quantile plot of hidden unit prior distributions *P*(*h*) between teacher and students, for the example teacher from Fig. 3 and its five students. (corresponds to column 3 of Fig. S5 A).

**Fig S7.**
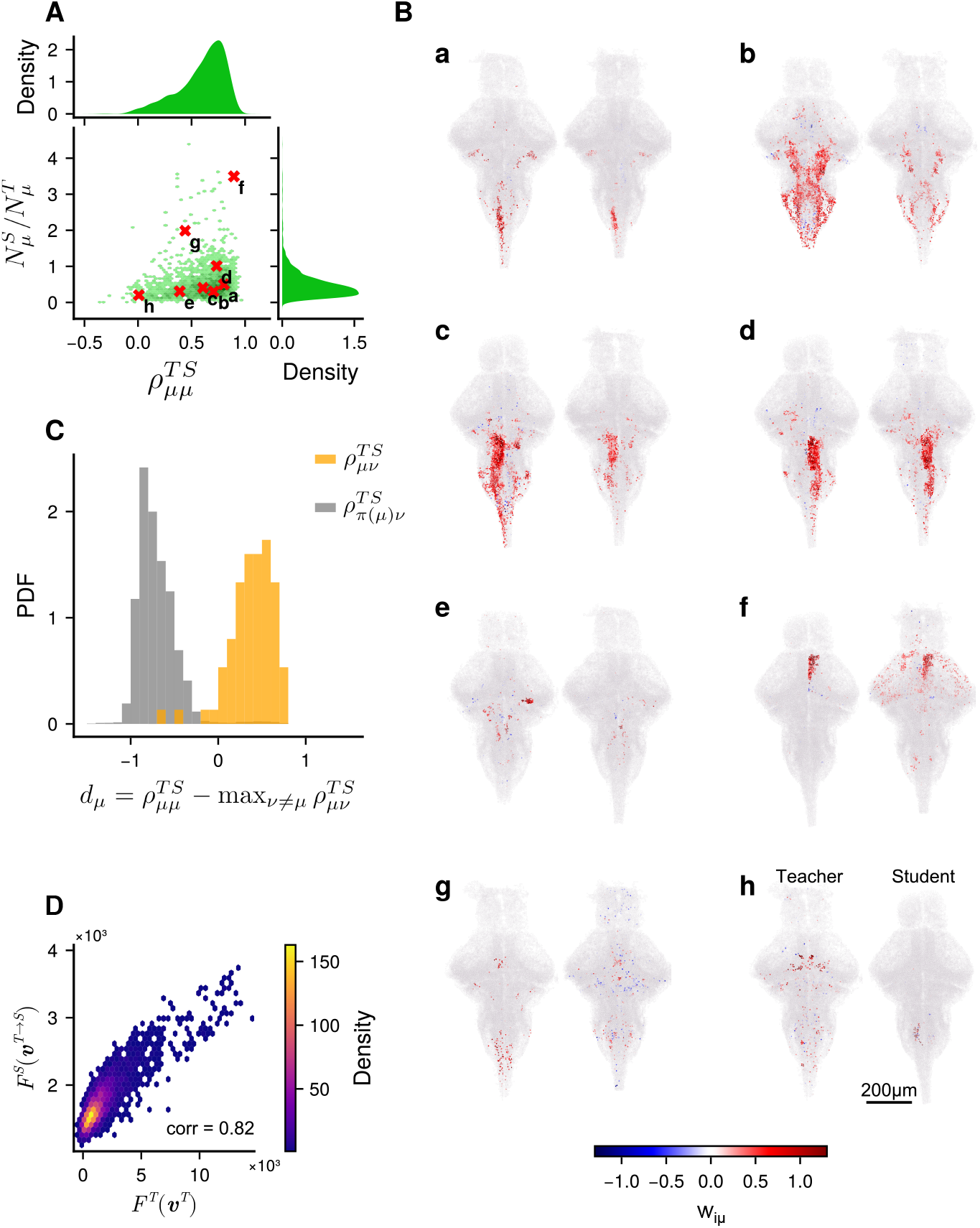
Supplementary for Fig. 3 on the measurement of distances between weight maps. **A**: Joint distribution for the spatial correlation 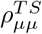 (see Methods Measuring Spatial Similarity) of weight maps between teacher and students, and ratio 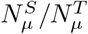 of the number of neurons 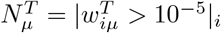 significantly connected with hidden unit µ in the teacher and in the student *N* ^*S*^. This distribution was created for all hidden units in all teacher-student pairs. Labeled red crosses correspond to the maps presented in panel B. **B**: Example hidden unit weight maps for teacher (left) and student (right) RBMs. Each map pair corresponds to a red cross in panel A. **C**: Distribution of 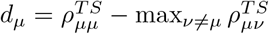 (see Methods Measuring Diagonal Dominance) for all hidden units µ of the example teacher-student pair from Fig. 3. In orange the distribution for the observed matrix 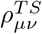, and in gray for the shuffled matrix 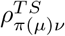 . **D**: Supplementary for Fig. 4. Free energy of an example teacher’s neuronal configurations, in the teacher RBM (*F* ^*T*^ (**v**^*T*^ )), versus translated in an example student RBM (*F* ^*S*^(**v**^*T* →*S*^)). Pearson correlation ∼ 0.82.

**Fig S8.**
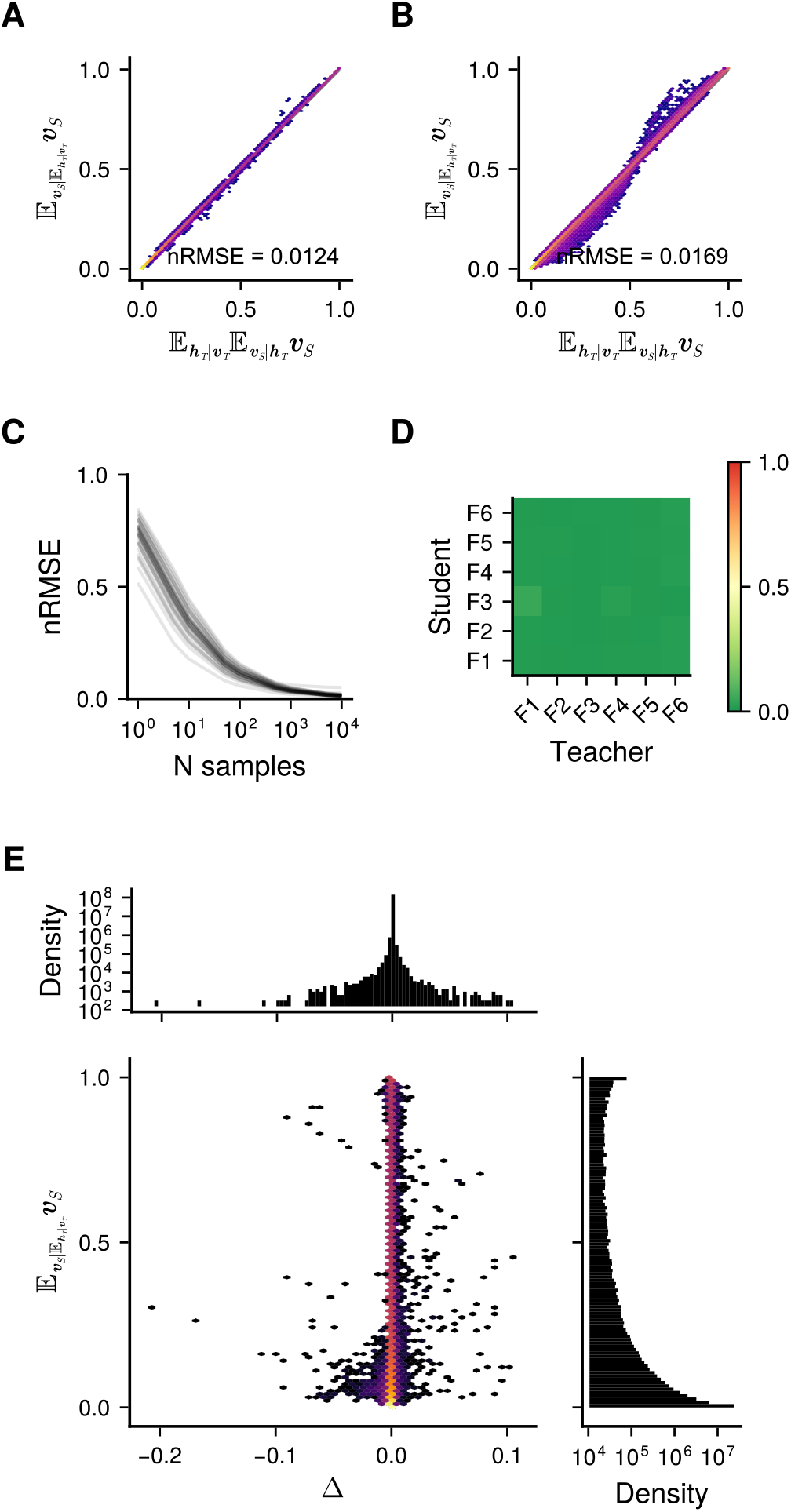
Supplementary for Fig. 4 and Methods Translating neuronal activity patterns across individuals on the method of translating neuronal configurations between fish. **A**: Identity plot between the two methods (sampling and expectation) activity transfer presented in Methods Translating neuronal activity patterns across individuals for an example teacher-student RBMs, and 10000 samples. **B**: Same as panel A for all pairs of teacher and student combined. **C**: nRMSE (see Methods Measuring identity) between the 2 methods (see panel A) for all teacher-student pair (one line per pair), as a function of the number of samples used. **D**: nRMSE for activity translation between every teacher-student pair (and activity reconstruction of teacher along the diagonal) for 10000 samples. **E**: Joint distribution of the sampling method and the corrective factor Δ.

**Fig S9.**
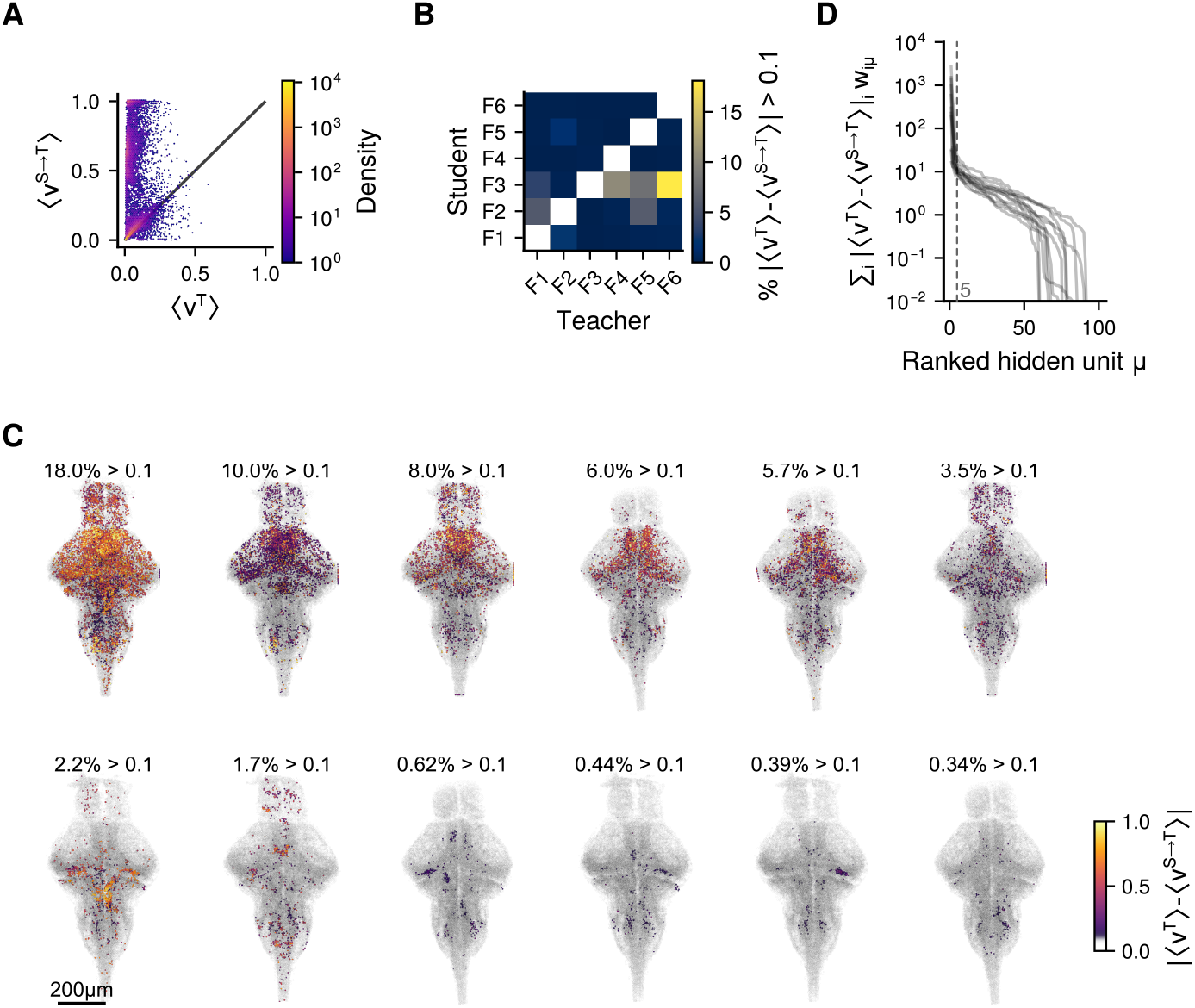
Supplementary figures for Fig. 4 on the outliers presented in Fig. 4 C,E,G A: Stereotypy of average temporal neuronal activity ⟨v_i_⟩ _*t*_ between an example teacher ( ⟨**v**^T^⟩ _*t*_), and an example student translated into the teacher ( ⟨**v**^S→*T*^⟩ _t_). We chose specifically the worst outlier. Neurons which lie far from the identity line are neurons whose mean activity are poorly reproduced when translating activity from one fish to the other. This per-neuron error is quantified as the translation residual 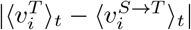. **B**: Percentage of neurons in each student with a translation residual greater than 0.1. **C**: Map of the translation residual per neuron, for the 12 worst teacher-student pairs. **D**: Translation residuals projected onto hidden units 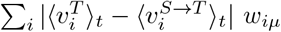. One line per teacher-student pair presented in panel C. Projected residuals are systematically dominated by less than 5 hidden units (vertical dashed line).

**Fig S10.**
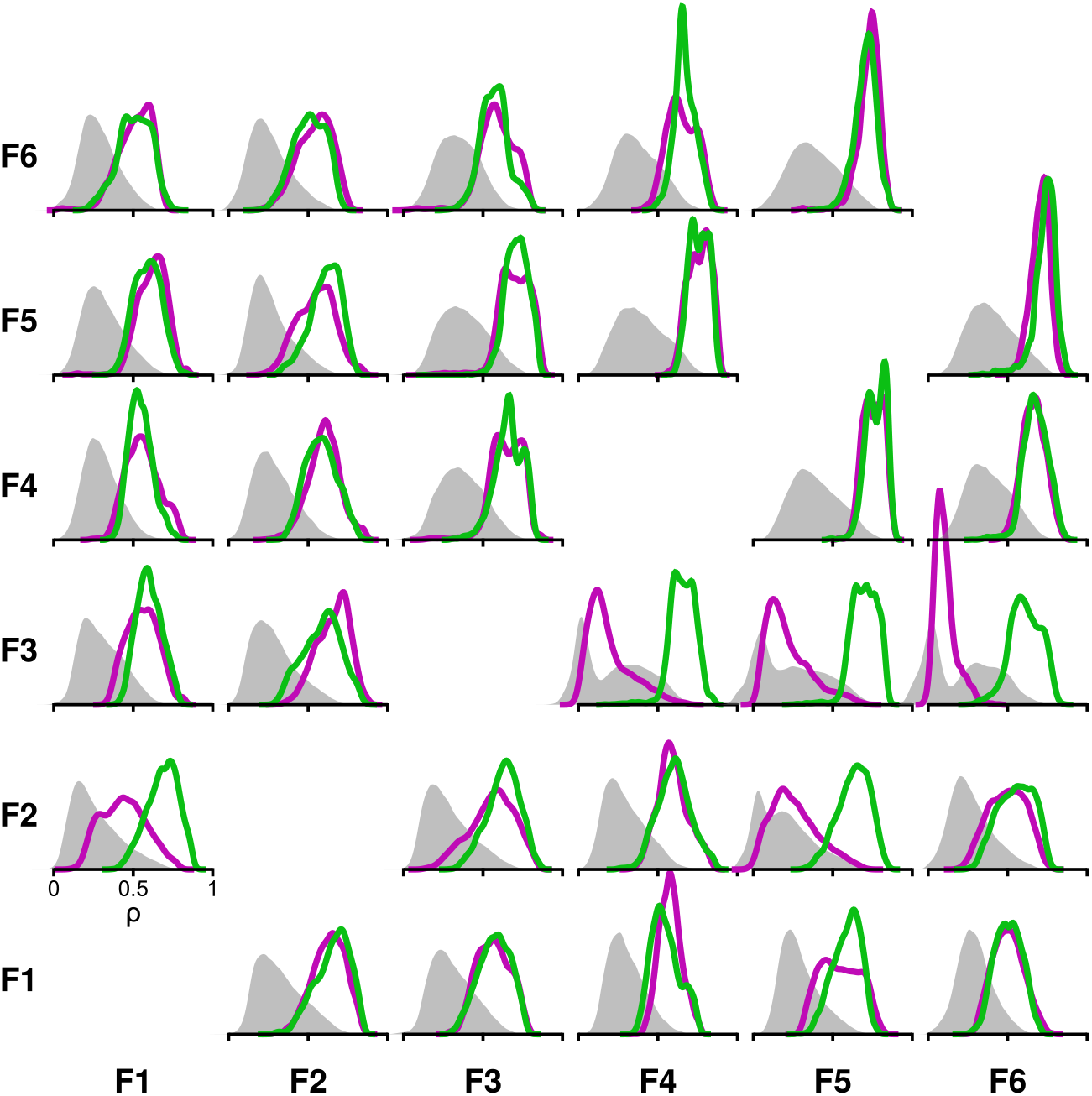
Supplementary figures for Fig. 4 of the measurement of distances between weight maps. Same as Fig. 4 F for every teacher-student pair. Spatial correlation 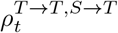 (magenta) and 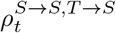 (green) between reconstructed and translated neuronal configuration for each time frame of an example teacher-student pair. In gray, the distribution of spatial correlations between shuffled pairs of time frames.

### Hyper-parameters selection

In this work, we used two sets of hyperparameters because the voxelized and single-cell RBMs operate at very different visible-layer dimensionalities and statistics. In both cases, the number of hidden units *M* and the regularization coefficient *λ* were chosen by cross-validation based on the model’s ability to reproduce the moments: ⟨v_i_⟩, ⟨h_µ_⟩, ⟨v_i_h_µ_⟩, ⟨v_*i*_*v*_*j*_⟩ − ⟨v_i_⟩⟨*v*_*j*_⟩ and ⟨h_µ_h_ν_ ⟩ − ⟨h_µ_⟩⟨*h*_*ν*_ ⟩, as described in Methods Evaluation.

#### Single-cell LaRBMs

We used *M* = 100 and *λ*_1_ = 0.02. This choice follows previous cross-validation performed on a single-fish dataset by van der Plas et al. [41], which found an optimum number of hidden units *M* = 200 for a fish with ∼55k segmented neurons. In this article, the authors also used *M* = 100 for fish with ∼30k neurons, maintaining a comparable hidden-to-visible ratio. To avoid repeating this computationally intensive cross-validation, to remain consistent with van der Plas et al. [41], and to reduce training duration, we chose *M* = 100 hidden units for our analyses. Importantly, because we use L1 weight regularization, increasing *M* does not necessarily yield a finer-grained representation as a substantial fraction of hidden units becomes effectively disconnected from the visible layer at high *M* (see added sensitivity analysis below).

#### Voxel-level RBMs

Here we used *M* = 40 hidden units because voxelization reduces the visible layer to ∼3k units and changes the correlation structure. For this case, cross-validation was performed explicitly in the present work (Methods Training; Fig. S3 A-B): we first carried out a broad grid search on one fish, then refined the search for the concatenated dataset of six fish.

### LaRBM sensitivity to changes in hyper-parameters

To directly assess sensitivity to *M* and λ_1_ for the LaRBMs, we performed an additional parameter sweep using the example teacher–student pair shown in the manuscript (Fig. S11). Across *M* ∈ [50, 200] and λ_1_ ∈ [0.005, 0.04], we found:

1. Teacher models converge and accurately reproduce the teacher fish statistics over a broad region of *M* and λ (Fig. S11 A).
2. As *M* and λ increase, the fraction of effectively disconnected hidden units increases (Fig. S11 B), showing that higher nominal *M* does not automatically translate into a larger effective latent representation under L1 regularization.
3. Student models trained by these teachers remain qualitatively stable for *M* ≥ 80. By contrast, insufficient regularization (λ_1_ < 0.02) leads to failures in reproducing key statistics (Fig. S11 C).
4. The free-energy distributions for reconstructed and translated patterns, the central energy-based comparison used in the manuscript, remain comparable throughout the stable region, with the main deviations again occurring for under-regularized models (Fig. S11 D).

Outside the range *M* ∈ [50, 200] and λ_1_∈ [0.005, 0.04], teacher-RBMs systematically failed to converge.

Overall, these results show that the qualitative conclusions emphasized in the manuscript (successful convergence of teacher/student RBMs, faithful modeling of moments, and comparable free-energy structure after translation) are robust to *M* across a wide range, while being more sensitive to insufficient regularization.

**Fig S11.**
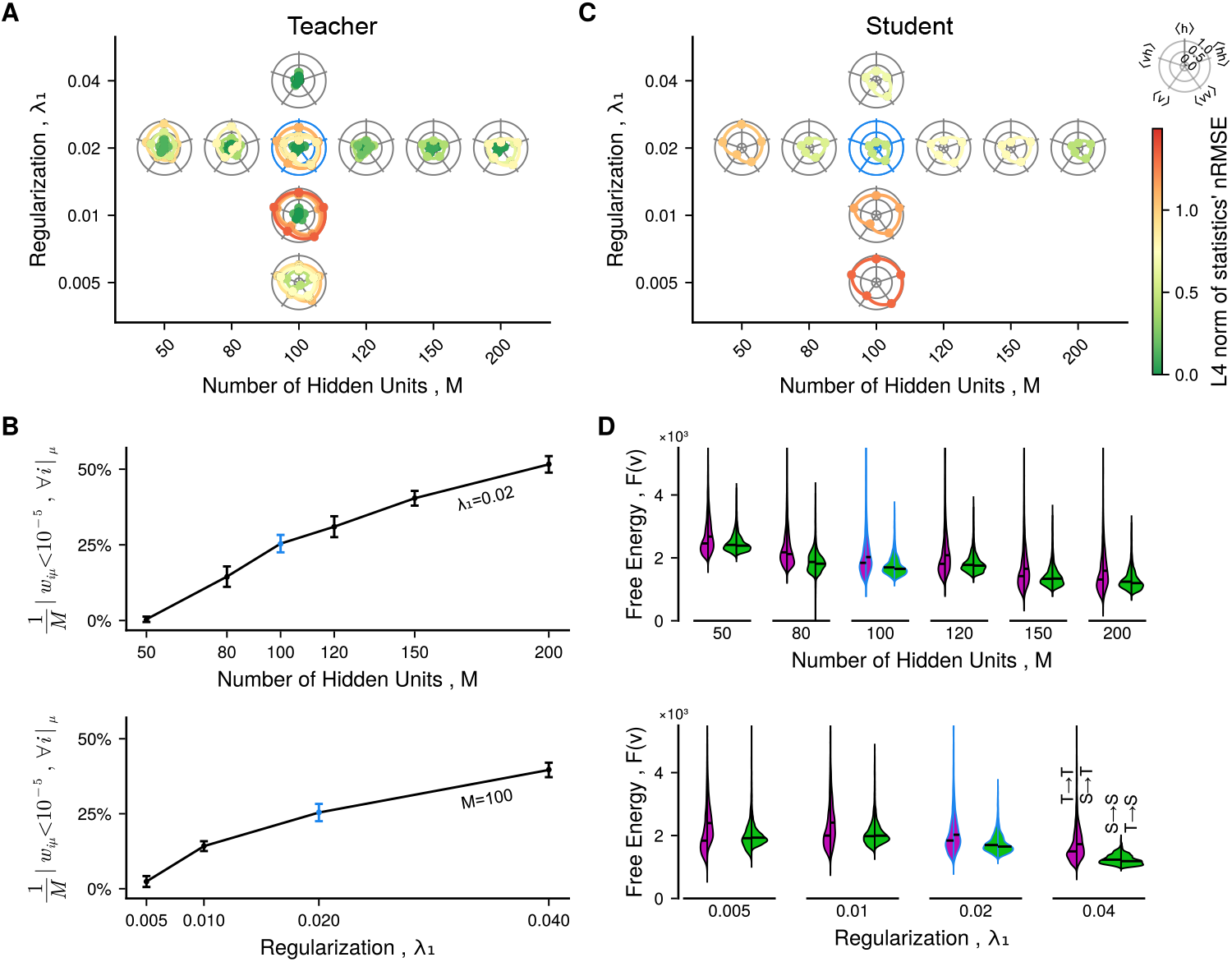
Supplementary figure for Section Hyper-parameters selection on the stability of the LaRBM to choices of hyperparameters. **A**: nRMSE (see Methods Measuring identity) of all five moments (see Methods Evaluation) used to evaluate RBM convergence on the example teacher fish. Each RBM is represented by a line in polar coordinates where every ray corresponds to a moment *m* ∈ {⟨**v**⟩, ⟨**vh**⟩, ⟨**h**⟩, ⟨**vv**⟩ − ⟨**v**⟩⟨**v**⟩, ⟨**hh**⟩ − ⟨**h**⟩⟨**h**⟩ }, and the ray length corresponds to the nRMSE_*m*_(data, generated) between empirical data and data generated by the RBM. Five training were performed and evaluated per hyperparameter pair (*M, λ*_1_), where *M* is the number of hidden units, and *λ*_1_ is the regularization factor (see Methods Training). **B**: Fraction of hidden units disconnected from the visible layer (*w*_*iµ*_ *<* 10^−5^ for all visible units *i*). Error bars represent mean and standard deviation computed across 5 RBM. Top: as function of the number of hidden units *M* and with a fixed regularization coefficient λ_1_ = 0.02. Bottom: as function of the regularization coefficient λ_1_ and for a fixed number of hidden units *M* = 100. **C**: Same as panel A, for the student LaRBM trained with the example teacher-student fish pair. **D**: Distributions of free energy. Left violin plots (purple) represent distributions computed in the teacher RBM, with reconstructed configurations (*T* →*T* ) on the left and translated configuration (*S* →*T* ) on the right. Right violin plots (green) represent distributions computed in the student LaRBM, with reconstructed configurations (*S* →*S*) on the left and translated configuration (*T* →*S*) on the right. Horizontal black lines indicate median. Top: as function of the number of hidden units *M* and with a fixed regularization coefficient λ_1_ = 0.02. Bottom: as function of the regularization coefficient λ_1_ and for a fixed number of hidden units *M* = 100. *Throughout this figure, items in blue correspond to the hyperparameters pair* (*M* = 100, λ_1_ = 0.02) *used in the rest of this work*.

### Small weight expansion of the LaRBM

Consider a student LaRBM, with the training objective:

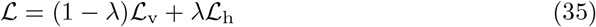

where 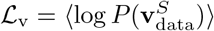 is the log-likelihood of neural activity under the student, 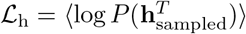 is the log-likelihood of hidden activity sampled from the teacher under the student, and λ ∈ [0, 1] is a tunable coefficient. Note that Eq. (35) is simply a rewrite of (2) in the main text.

In this section we consider a second-order expansion of this training objective (35), around small weights. Our goal is to gain some insight on how the trained weights are affected by the combination of the two training objectives, ℒ_v_ and ℒ_h_. For simplicity, we will assume that the units have been standardized to have zero means and unit variances.

The expansion of ℒ_v_ mirrors the expansion of ℒ_h_, after transposing the weight matrix. Therefore, we will first consider the expansion of ℒ_v_ in detail. We begin by defining the cumulant generating function of the hidden unit activities,

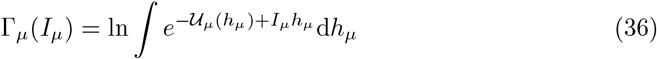

where *I*_*µ*_ = *I*_*µ*_(**v**) =∑ _*i*_ *w*_*iµ*_*v*_*i*_ are the inputs from the visible to the hidden layer.

Then, the free energy of the visible activity, after marginalizing over the hidden unit configurations, writes:

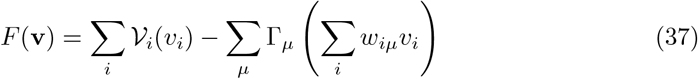

The function Γ_*µ*_(*I*_*µ*_) is called the cumulant generating function, because the coefficients appearing in its Taylor series are the cumulants of the hidden unit activities at zero weights:

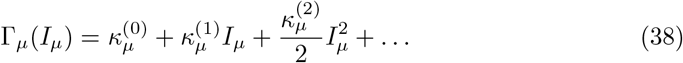

where 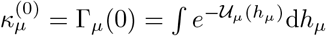, while 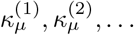 are the cumulants of the hidden unit distributions of different orders, at zero weights. For the first few orders, these are the mean, variance, and so on:

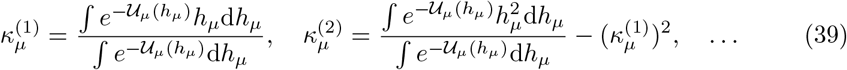

It follows that we can expand the free energy *F*(**v**) in powers of the weights around the origin as:

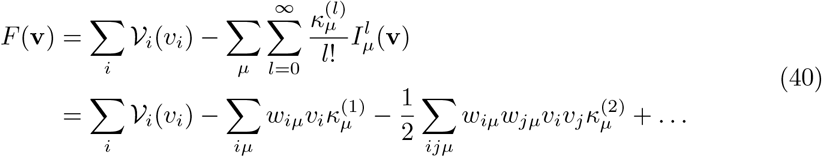

Now let’s look at the partition function,

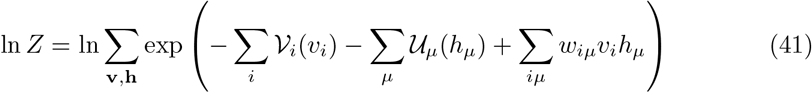

We begin by denoting the measure at zero weights by:

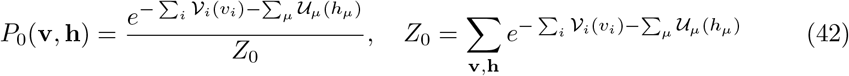

Then we can write:

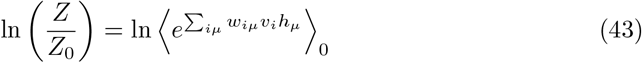

where ⟨·⟩_0_ denotes the average under *P*_0_. We can treat this expression as a cumulant generating function, which then allows us to write the small weight expansion as follows:

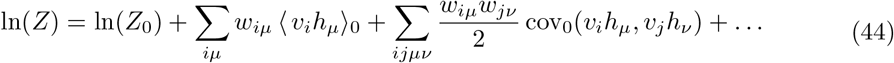

where the averages and covariances are under *P*_0_.

We can now substitute the previous equations to obtain the expansion of ℒ_v_ as follows:

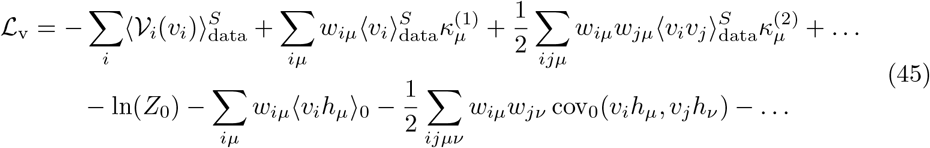

We will assume for simplicity that the hidden units are initialized such that 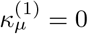 and 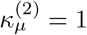. Then at zero weights it follows that ⟨v_i_ h_*µ*_ ⟩_0_ = 0. Moreover,

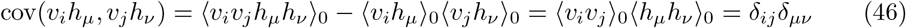

because different units are independent at zero weights. Therefore, omitting terms that are independent of the weights, we arrive at the simplified expansion:

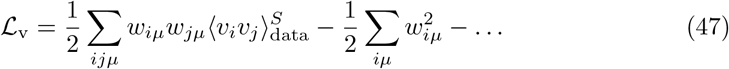

Mirroring this derivation, we obtain a similar expansion for ℒ_h_:

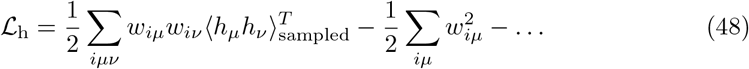

where we assume that the visible units are initialized with zero means and unit variances, for simplicity.

Combining the two:

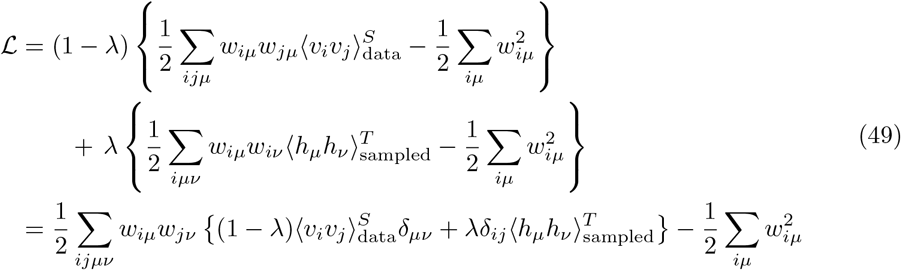

We now try to understand the eigenmode structure of the matrix 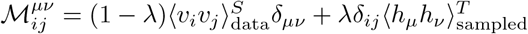. If *x*_*iµ*_ is an eigenvector with the eigenvalue τ,

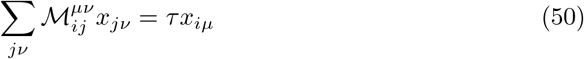

or

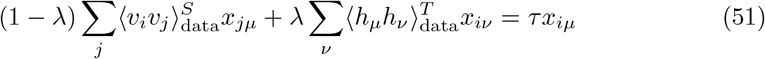

We search for eigenvectors of the factorized form *x*_*iµ*_ = *x*_*i*_*x*_*µ*_. Substituting:

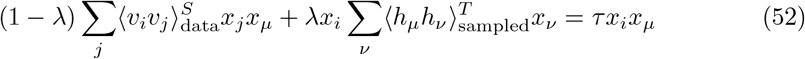

We then assume that *x*_*i*_ is an eigenvector of 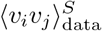, with eigenvalue τ_v_, and that *x*_*µ*_ is an eigenvector of 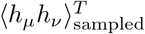 with eigenvalue τ_h_. Then,

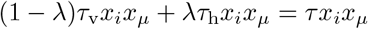

or (1 − λ)τ_v_ + λτ_h_ = τ . Since 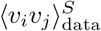 has *N* eigenvectors and 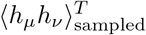 has *M* eigenvectors, this construction yields all *NM* eigenvectors of 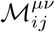.

This derivation allows us to offer some insight into why the student LaRBM weights often have smaller magnitude than the weights of the teacher RBM. We find empirically that the eigenvalues of 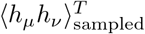 are often of lower magnitude than the eigenvalues of the covariance matrix of the data used to train the student, 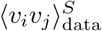 (see Fig. S12). This implies that the histogram of eigenvalues of the Hessian matrix of the combined objective ℒ for the student will be shifted to lower values. The corresponding directions in weight space are then more strongly suppressed by the regularization, and will thus be lower.

**Fig S12.**
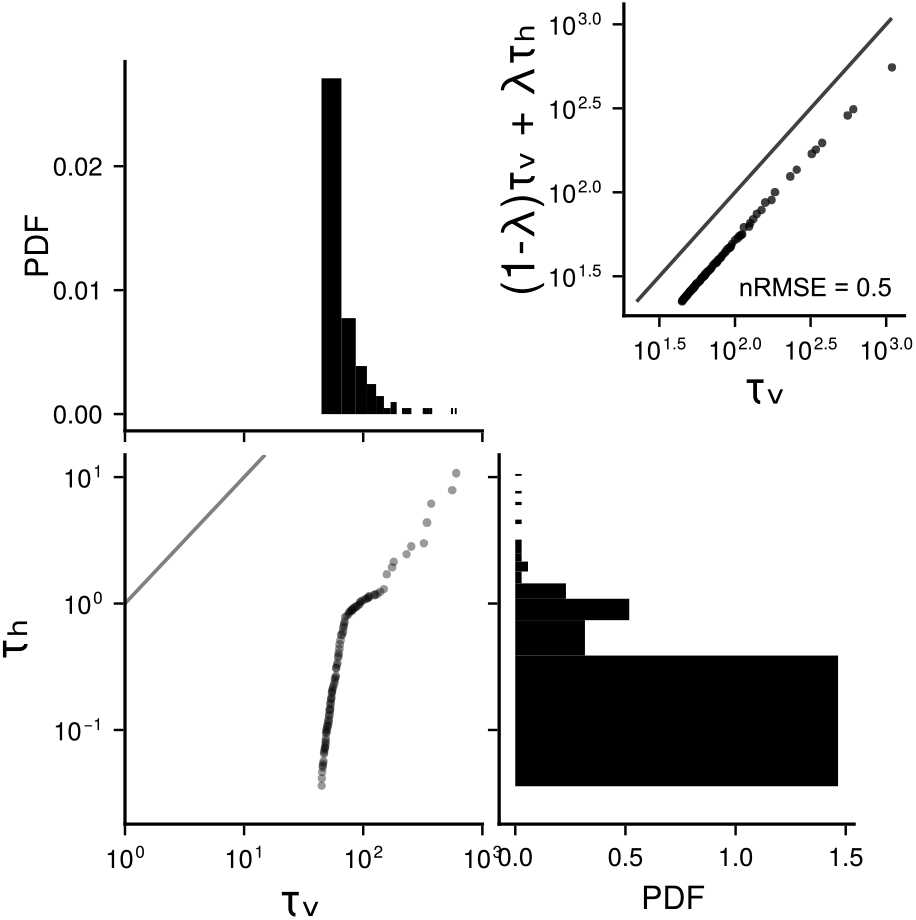
Supplementary figures for Section Small weight expansion of the LaRBM on the eigenvalues of of the LaRBM training objective. Top left: histogram of the top 99 eigen values τ_v_ of the correlation matrix 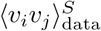 of visible data in the student. Bottom right: histogram of the top 99 eigen values τ_h_ of the correlation matrix 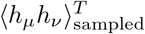 of hidden configuration generated from the teacher RBM. Bottom left: Joint scatter of the τ_v_ and τ_h_ from the previous two panels. Top Right: QQ plots of the distributions τ_v_ and (1 − λ)τ_v_ + λτ_h_ with λ = 0.5.

